# Spatial proteogenomics reveals distinct and evolutionarily-conserved hepatic macrophage niches

**DOI:** 10.1101/2021.10.15.464432

**Authors:** Martin Guilliams, Johnny Bonnardel, Birthe Haest, Bart Vanderborght, Anna Bujko, Liesbet Martens, Tinne Thoné, Robin Browaeys, Federico F. De Ponti, Anneleen Remmerie, Camille Wagner, Bavo Vanneste, Christian Zwicker, Tineke Vanhalewyn, Amanda Gonçalves, Saskia Lippens, Bert Devriendt, Eric Cox, Giuliano Ferrero, Valerie Wittamer, Andy Willaert, Suzanne J.F. Kaptein, Johan Neyts, Kai Dallmeier, Peter Geldhof, Stijn Casaert, Bart Deplancke, Peter ten Dijke, Anne Hoorens, Aude Vanlander, Frederik Berrevoet, Yves Van Nieuwenhove, Yvan Saeys, Wouter Saelens, Hans Van Vlierberghe, Lindsey Devisscher, Charlotte L. Scott

**Author notes:** These authors contributed equally (Martin Guilliams, Johnny Bonnardel, Birthe Haest, Bart Vanderborght, Wouter Saelens, Hans Van Vlierberghe, Lindsey Devisscher, Charlotte L. Scott). These authors contributed equally.

## Abstract

The liver is the largest solid organ in the body, yet it remains incompletely characterized. Here, we present a spatial proteogenomic atlas of the healthy human and murine liver combining single-cell CITE-seq, single-nuclei sequencing, spatial transcriptomics and spatial proteomics. By integrating these multi-omic datasets, we provide validated strategies to reliably discriminate and localize all hepatic cells. We then align this atlas across seven species, revealing the conserved program of *bona fide* Kupffer cells and bile-duct macrophages. We also uncover the respective spatially-resolved cellular niches of these macrophages and the microenvironmental circuits driving their unique transcriptomic identities. We demonstrate that bile-duct macrophages are induced by local lipid exposure, while Kupffer cells crucially depend on their crosstalk with hepatic stellate cells via the evolutionarily-conserved ALK1-BMP9/10 axis.

## Introduction

The immense technological advances in single-cell transcriptomics have enabled a better understanding of the cellular composition of different organs across species. However, we still lack information regarding how these cells are organized in their distinct microenvironmental niches. Moreover, the specific cell-cell interactions determining the identity of individual cells within tissues remain to be defined (Guilliams and Scott, 2017; Lindeboom et al., 2021). While the spatial organization of hepatocytes within the liver is understood (Halpern et al., 2017), that of non-parenchymal liver cells remains unclear. This is the case for the mouse liver, but even more so for the human liver, where the identity and the precise localization of most hepatic cells is unknown. Moreover, the link between the transcriptome and the proteome has not been studied, resulting in a lack of reliable surface markers to identify, purify or localize these cells by flow cytometry and confocal microscopy. Here, we have used proteogenomics techniques including CITE-seq and spatial approaches to identify all cells and their specific locations within the healthy livers of mice and humans. By doing so, we have developed strategies for the identification and further study of the different cell types. Demonstrating the usefulness of this approach, with this information, we also identify the conserved spatially-relevant signals driving distinct hepatic macrophage phenotypes in the healthy liver.

## Results

### A practical proteogenomic atlas of the murine liver

To generate a proteogenomic atlas of the liver, we first examined the optimal method for retrieving all hepatic cells. Using the murine liver, we compared single-cell RNA-sequencing (scRNA-seq) using cells isolated via *ex vivo* or *in vivo* enzymatic digestion with single-nuclei RNA-sequencing (snRNA-seq). With each technique we observed a distinct cellular composition (Fig. S1A-C). While snRNA-seq yielded a lower number of genes/cell than scRNA-seq, it best recapitulated the cell frequencies observed *in vivo* (Fig. S1D-L). Given the distinct cellular compositions with each method, to ensure all cells could be profiled, we opted to use a combination of all protocols in our study. To investigate mRNA and protein expression at single cell resolution, we used cellular indexing of transcriptomes and epitomes by sequencing (CITE-seq) (Stoeckius et al., 2017). Thus, we stained a selection of the scRNA-seq samples with 107-161 oligo-conjugated antibodies (Fig. 1A). Data were pooled together for a single analysis where, with TotalVI (Gayoso et al., 2019), both the protein and mRNA profiles were considered for clustering (Fig. 1B). Analysis of the differentially expressed genes (DEGs; Fig. S2A & Table S1) and proteins (DEPs; Fig. S2B & Table S2) identified 17 cell types (Figs. 1B, S2C). Moreover, we identified surface markers for all cells, including VSIG4 and FOLR2 for Kupffer cells (KCs) (Fig. S2B,D).

**Figure 1:**
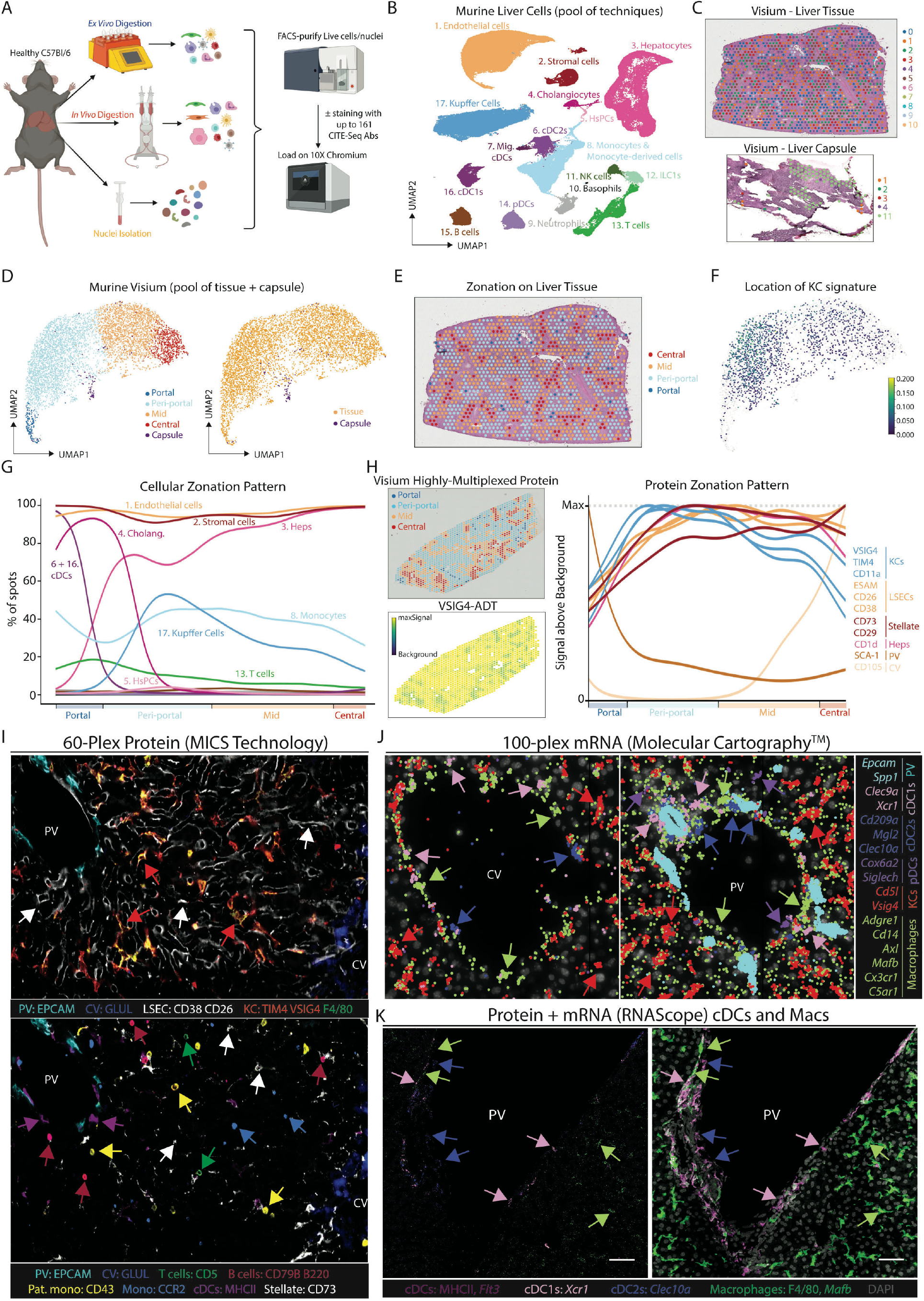
A proteogenomic atlas of the healthy murine liver. (A) Hepatic cells were isolated from healthy C57B/l6 mice by *ex vivo* (5 mice, 15 samples) or *in vivo* (5 mice, 19 samples) enzymatic digestion. Alternatively, livers were snap frozen and nuclei isolated by tissue homogenization (4 mice, 12 samples). Live cells/intact nuclei were purified using FACS. For cells, total live, live CD45^+^, live CD45^−^, live hepatocytes or myeloid cells (live CD45^+^, CD3^−^, CD19^−^, B220^−^, NK1.1^−^) were sorted. 18 cell preparations (7 *ex vivo,* 11 *in vivo)* were also stained with a panel of 107-161 barcode-labelled antibodies for CITE-seq analysis. All datasets were pooled together and after QC 185894 cells/nuclei were clustered using TotalVI. (B) UMAP of sc/snRNA-seq data. (C) Tissue and capsule images from Visium analysis with clusters overlaid. (D) UMAP of zonation of Visium spots (left) & origin of the cells (right). (E) Zonation pattern mapped onto tissue slice. (F,G) Indicated cell signatures from sc/snRNA-seq mapped onto the Visium zonation data. (H) mRNA zonation pattern in Visium Highly-Multiplexed Protein analysis and VSIG4-ADT expression pattern (left) and zonated expression patterns of indicated antibodies (right). (I) MICS analysis of indicated proteins and cell types. (J) Molecular Cartography of indicated genes and cell types. (K) mRNA (*Xcr1, Flt3l, Mafb* and *Clec10a*) and protein (MHCII and F4/80) expression in the same tissue slice. Scale bar 50μm. PV; portal vein, CV; central vein. Arrows indicate specific cell types, where color corresponds to cell type/markers. All images are representative of 2-4 mice.

### Distinct spatial orientation of hepatic myeloid cell subsets

To locate the cells identified we performed spatial transcriptomics analysis using Visium. For this, we cut the liver in two distinct orientations to profile both the liver tissue and the capsule (Figs. 1C, S2E). We ordered each Visium spot along a spatial trajectory, and annotated portal, periportal, mid and central zones based on known hepatocyte zonation markers (Halpern et al., 2017) (Figs. 1D,E, S2F). By using the reference sc/snRNA-seq data, we then deconvolved each spot into its constituent cell types and investigated how cell abundance changed with zonation (Figs. 1F,G, S2G). Validating this approach, as reported, cholangiocytes mapped specifically to the portal zones (Aizarani et al., 2019), while KCs were located in peri-portal and mid zones (Bonnardel et al., 2019; Gola et al., 2021). Moreover, we identified T cells, endothelial cells (ECs) and stromal cells (SCs) across all zones, while conventional dendritic cells (cDCs) were found at the portal vein, with a minor presence at the central vein (Figs. 1F,G, S2G).

To validate these locations at single-cell resolution, we next sought to identify the best cell-specific surface markers that would also work by confocal microscopy. As the fixation step utilized for confocal microscopy often affects the integrity of different epitopes, it is not possible to predict which antibodies that work on single cell suspensions will work spatially on fixed and intact tissue. Therefore, to simultaneously screen multiple antibodies to identify those working by microscopy, we performed a second Visium analysis which we complemented with 100 oligo-conjugated antibodies (Visium highly-multiplexed protein), chosen based on the CITE-seq results (Fig.1H). The antibodies identified to work spatially were then validated at single-cell resolution, using MACSima™ Imaging Cyclic Staining (MICS) technology and a 60-plex antibody panel (Fig. 1I). Unfortunately, we could not identify useful surface markers for all populations, for example we did not identify enough discriminatory surface markers that worked by confocal microscopy to distinguish the cDC subsets spatially. Thus, to confirm the locations of these cell subsets we turned to Molecular Cartography™ (Resolve BioSciences) that allows for 100-plex spatial mRNA analysis. Genes were selected based on the DEGs from the sc/snRNA-seq data that were also spatially-resolved according to Visium. We also identified the portal-central trajectory in this dataset using cholangiocyte genes (*Epcam, Spp1*) and known zonated hepatocyte genes (*Glul, Cyp2e1, Hal, Sds*; Fig. S2H). Using expression of *Xcr1, Clec9a* (cDC1s) and *Cd209a, Mgl2* and *Clec10a* (cDC2s), we confirmed that both cDC1s and cDC2s were localized primarily at the portal vein (Fig. 1J). As cDC2s shared a number of genes with monocyte-derived cells (Fig. S2A), we also examined the expression of general monocyte/macrophage (*Cd14, Adgre1, Axl, Mafb, Cx3cr1, C5ar1*) and KC-specific genes (*Cd5l, Vsig4*) to further validate their identification as *bona fide* cDC2s. This analysis further suggested that the cDC2s we identified were *bona fide* cDC2s, lacking any macrophage/monocyte markers, however, it also identified populations of macrophages at the portal and central veins distinct from both KCs and cDC2s (Fig. 1J). The punctate nature of mRNA expression in these analyses combined with the dendritic shape of myeloid cells renders it difficult to convincingly determine cell boundaries and to conclude these cDC2s and macrophages were distinct cells. To validate this, we therefore developed a protocol that combines mRNA detection (RNAScope) with surface protein detection. Examining expression of cDC- or macrophage-specific mRNAs combined with protein surface markers confirmed the presence of portal vein cDC1, cDC2s and non-KC macrophages (Fig. 1K). Taken together, by combining multiple spatial transcriptomic and proteomic approaches, we located all the cells within the murine liver and identified additional heterogeneity within the myeloid cells, not revealed when examining the sc/sn-RNA-seq dataset in isolation. This highlights the power of combining single-cell and spatial proteogenomic techniques to investigate cellular heterogeneity.

### Refined analysis of myeloid cells identifies three subsets of hepatic macrophages

To better understand these non-KC macrophages, we zoomed in on myeloid cells (cDCs, KCs, monocytes and monocyte-derived cells) in our sc/snRNA-seq analysis defining 11 populations (Fig. 2A-C & Tables S3,4). This included KCs, 3 populations of non-KC macrophages and cells that had a profile intermediate between monocytes and patrolling monocytes or macrophages, termed transitioning monocytes. Closer inspection of the non-KC macrophages identified cluster6 as peritoneal macrophages (Fig. 2B). The DEGs between the remaining populations suggested that cluster7 likely resembles capsule macrophages (Sierro et al., 2017), expressing *Cd207 and Cx3cr1* while cluster8 resembles *Gpnmb^+^Spp1*^+^ lipid associated macrophages (LAMs) we recently described in the fatty liver (Remmerie et al., 2020) (Fig. 2A-C). Conversion of the CITE-seq data into a flow cytometry file allowed an *in-silico* gating strategy to be defined (Fig. S3A). Validating this, we utilized the strategy to FACS-purify the populations and assess gene expression (Fig. S3B-D). Washing the liver prior to digestion enriched the peritoneal macrophages in the wash fraction, demonstrating these were contaminants on the liver surface rather than being present in the liver tissue itself (Fig. S3E). While the CITE-seq markers did not discriminate between clusters7 and 8, adding CD207 to the panel enabled the non-KCs to be divided into CD207^+^ and CD207^−^ macrophages (Fig. S3F). Fitting with their designation as capsule macrophages, the relative abundance of CD207^+^ macrophages was increased if we dissected and digested the capsule (Fig. S3F). However, although Molecular Cartography confirmed the presence of *Cd207^+^* macrophages in the capsule, it also revealed *Cd207^+^* macrophages at the central vein, which were rarely found at the portal vein (Figs. 2E-H & S3G-J). Thus, cluster7 consists of both capsule and central vein CD207^+^ macrophages. This finding further demonstrates the need for spatial approaches to confirm cell identities, as this signature was previously considered to be unique to the capsule macrophage population (Sierro et al., 2017). Molecular Cartography also identified macrophages at the portal and central veins expressing *Ccr2* and *Chil3* (Figs. 2G,H & S3H,J), resembling transitioning monocytes (cluster11). Finally, a population of *Gpnmb*-expressing macrophages were found to be specifically located around the bile-ducts (Fig. 2G,H & S3H,K). As *Gpnmb* expression is cluster8-specific (Fig. 2B) and these cells resemble LAMs (Remmerie et al., 2020), we termed these cells bile-duct LAMs.

**Figure 2:**
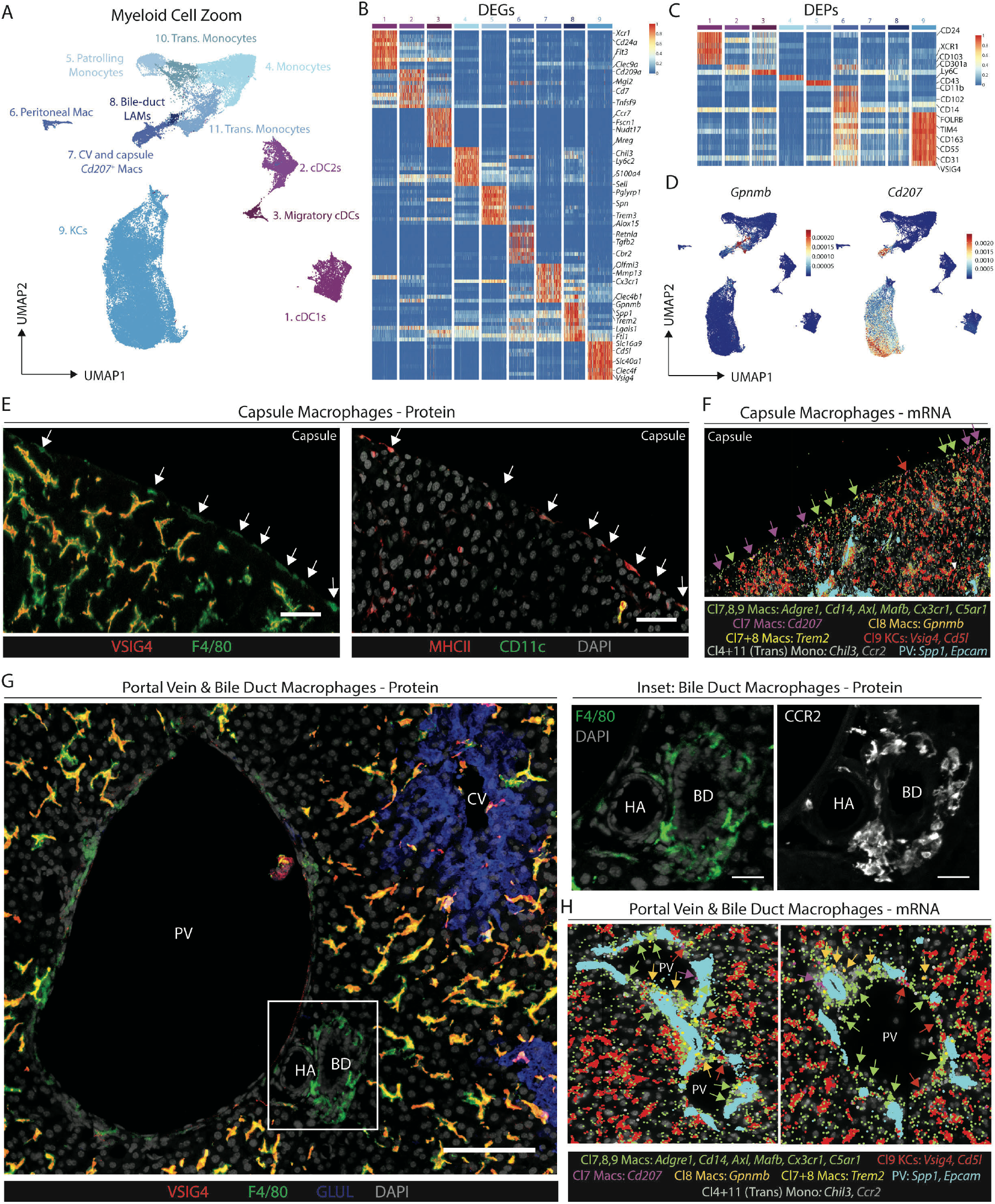
A population of macrophages reside around the bile-duct in the healthy murine liver. (A) UMAP of murine myeloid cells (71261 cells/nuclei) isolated from Fig. 1B and re-clustered with TotalVI. (B,C) Top DEGs (B) and DEPs (C) between cell types identified. (D) Expression of *Gpnmb* and *Cd207*. (E) Expression of VSIG4 and F4/80 (left) or MHCII, CD11c and DAPI (right) by confocal microscopy. Capsule macrophages (left) or MHCII^+^ capsule macrophages (right) identified by white arrows. Scale bar 50μm. (F) Molecular Cartography of indicated genes and cell types at liver capsule. (G) Expression of VSIG4, F4/80, GLUL and DAPI (left) or F4/80 or CCR2 (right, inset) by confocal microscopy. Scale bar 100μm. (H) Molecular Cartography of indicated genes and cell types at portal triad. PV; portal vein, CV; central vein, HA; hepatic artery and BD; bile duct. Arrows indicate specific cell types, where color corresponds to cell type/markers. All images are representative of 2-4 mice.

### Macrophage subsets reside in distinct spatial niches

As all the macrophage populations are in close contact with CD45^−^ cells in their local environment (Bonnardel et al., 2019) (Fig. S3L), we further analyzed the CD45^−^ cells, identifying multiple subsets of ECs and SCs and a gating strategy to distinguish them (Figs. 3A-C, S4A-C & Tables S5,6). ECs could be further subdivided into 4 distinct clusters and analysis of their locations allowed them to be identified as central vein ECs (cluster10), LSECs (cluster9), portal vein ECs (cluster11) and Lymphatic ECs (LECs; cluster12) (Figs. 3D,E, S4D,E). As Visium found fibroblasts at both the portal and central veins (Fig. 3D), and as a previous report has suggested the presence of distinct subsets within these cells (Dobie et al., 2019), we further zoomed in on the SCs to better assess their heterogeneity (Figs. 3F,G, S4A-C,F & Table S7). This revealed subsets of mesothelial cells and fibroblasts restricted to the capsule (Fig. 3H,I). *Myh11*^+^ vascular smooth muscle cells (VSMCs) were localized around hepatic arteries, portal veins and central veins (Figs. 3F-H,J & S4G) and *Mfap4^+^Svep1^+^Clic5^−^Reln^−^* fibroblasts were found to be central vein fibroblasts (Figs. 3G-H,J, S4G). Finally, we identified a subset of *Clic5^+^Reln^+^* fibroblasts (cluster3) which were localized around the cholangiocytes and that we termed bile-duct fibroblasts (Figs. 3J & S4G). Taken together, the presence of these spatially-distinct subsets of ECs and SCs highlights the uniqueness of the specific microenvironments in which the distinct macrophage populations reside.

**Figure 3:**
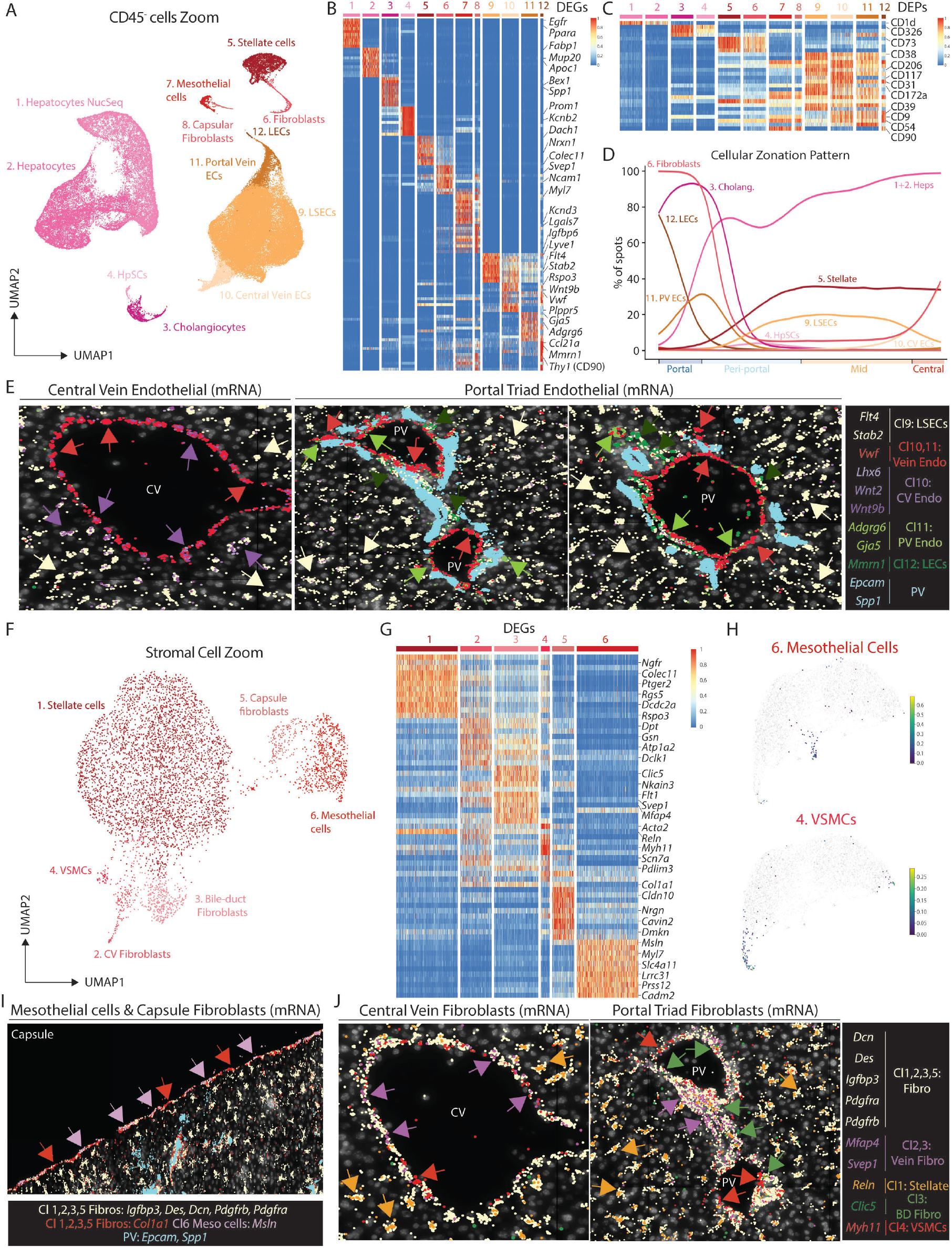
Hepatic macrophage populations reside in distinct niches. (A) UMAP of murine CD45^−^ cells (83410 cells/nuclei) isolated from Fig. 1B and re-clustered with TotalVI. (B,C) Top DEGs (B) and DEPs (C) between cell types identified. (D) Indicated cell signatures from sc/snRNA-seq mapped onto the Visium zonation data. (E) Molecular Cartography of indicated genes and cell types at central vein (left) and 2 different portal triads (center and right). (F) UMAP of murine stromal cells (5430 cells/nuclei) isolated from the UMAP in Fig. 3A and re-clustered with scVI. (G) Top DEGs between different cell types identified. (H) Identification of Mesothelial cell (top) and VSMC (bottom) signatures on zonated Visium data. (I) Molecular Cartography of indicated genes and cell types at the liver capsule. (J) Molecular Cartography of indicated genes and cell types at the central vein (left) and portal triad (right). PV; portal vein, CV; central vein, HA; hepatic artery and BD; bile duct. Arrows indicate specific cell types, where color corresponds to cell type/markers. All images are representative of 2-4 mice.

### A practical proteogenomic atlas of the healthy human liver

To determine the degree of conservation between the macrophage subsets and their different microenvironmental niches between the mouse and the human liver, we next generated a proteogenomic atlas of the human liver using sc/snRNA-seq and CITE-seq on 19 liver biopsies (Figs. 4A,B, S5A,B & Tables S9,10). Of these, most were histologically healthy with only 5 patients showing >10% hepatic steatosis (Table S8). Cellular proportions varied according to the isolation technique used, and while there was some variability between patients, this was not linked to the surgery (Fig. S5C-E). As Visium reliably located murine hepatic cells, we used this to locate the cells of the human liver in 4 biopsies (Fig. 4C). As patients with >10% steatosis clustered separately from the healthy samples (<10% steatosis; Figs. 4D,E, S5F,G), we used the healthy samples to calculate a baseline zonation and then transferred this trajectory onto the steatotic samples (Figs. 4F, S5F,G). This identified the steatosis to be predominantly peri-centrally located in these patients (Fig. 4G). This fits with previous clinical studies demonstrating peri-central steatosis to be most common in NAFLD patients, especially in early disease where peri-portal areas are often spared (Chalasani et al., 2008; Kleiner and Makhlouf, 2016). Notably, the overall cellular distribution was not impacted by the presence of peri-central steatosis, although neutrophils were preferentially localized in steatotic zones (Fig. 4G).

**Figure 4:**
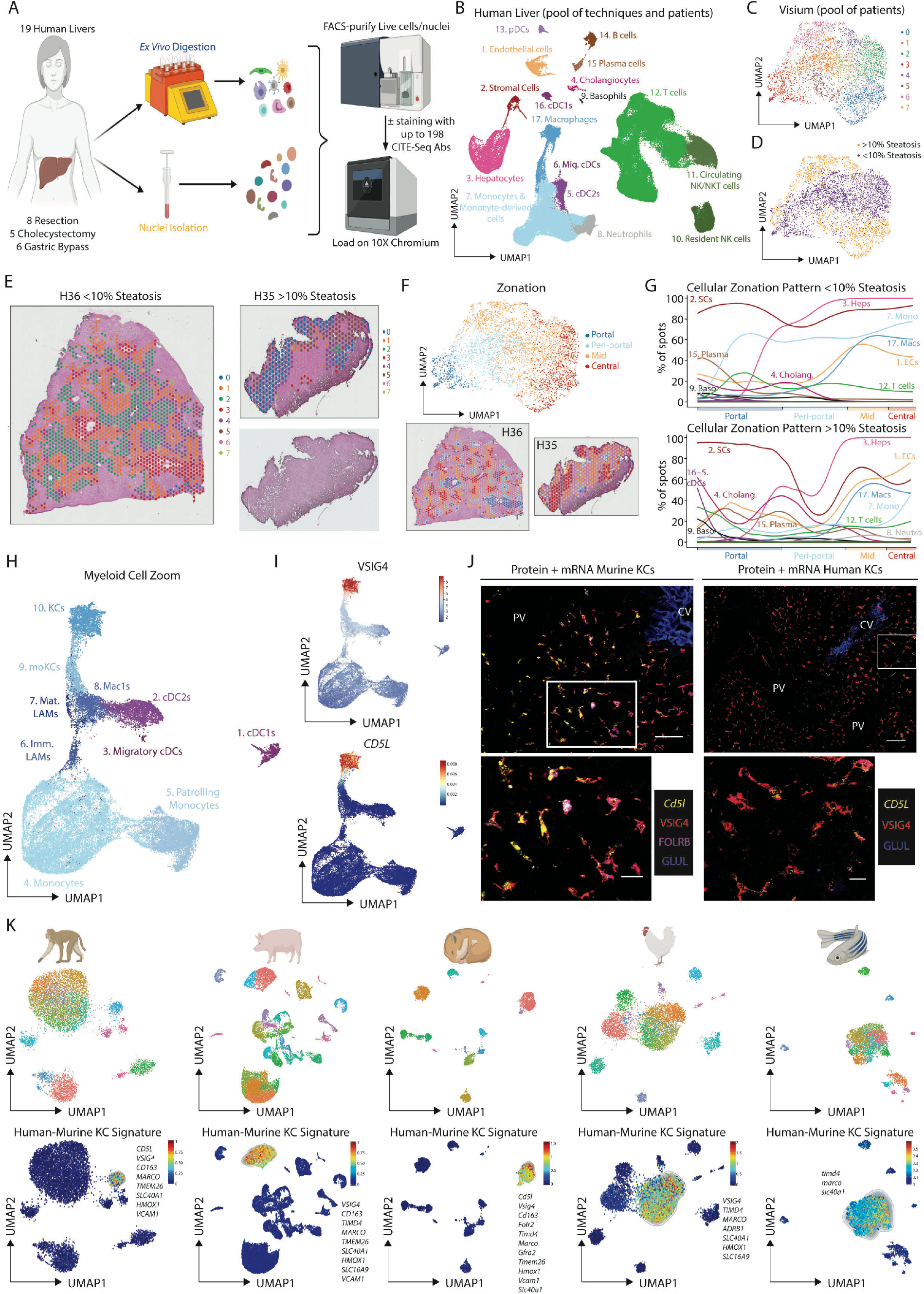
Identification of *bona fide* Kupffer cells across species. (A) Cells or nuclei were isolated from liver biopsies (∼1-2mm^3^; 14 cells, 5 nuclei) from patients undergoing either liver resection, cholecystectomy or gastric bypass. Live cells/intact nuclei were purified using FACS. Either total live, live CD45^+^, live CD45^−^ or lineage^−^cells (live CD45^+^,CD3^−^, CD19^−^) were sorted. 7 cell samples were stained with a panel of 198 barcode-labelled antibodies for CITE-seq analysis. All datasets were pooled together and after QC 167598 cells/nuclei were analyzed using TotalVI. (B) UMAP of sc/snRNA-seq data. (C) UMAP of Visium data from 4 patient biopsy samples. (D) Split of Visium spots based on %steatosis. (E) Healthy and steatotic Visium liver tissue with clusters overlaid and H+E staining to identify steatotic zones. (F) Zonation of Visium data (top) with zonation pattern mapped onto liver tissue (bottom). (G) Indicated cell signatures from sc/snRNA-seq mapped onto Visium zonation trajectory, healthy (top), steatotic (bottom). (H) Myeloid cells (40821 cells) were isolated from Fig. 4B and re-clustered with TotalVI. (I) Expression of VSIG4 protein (top) and *CD5L* mRNA (bottom). (J) Expression of VSIG4, F4/80, FOLRB, GLUL combined with *Cd5l/CD5L* on murine (left) and human (H25; right) livers. Scale bar 50μm. Inset in bottom panels. Scale bar 20μm. Images are representative of 2-4 mice/patients. (K) Livers (2/species) were isolated from healthy Macaque, Pig, Chicken, Hamster and Zebrafish. Cells were isolated by *ex vivo* digestion for CITE-seq (pig; 198 human antibodies) or scRNA-seq (hamster, chicken zebrafish) or Nuclei were isolated for snRNA-seq (macaque). Total live cells (Hamster, chicken, pig), DsRed^+^GFP^+^ cells (Zebrafish) or nuclei (macaque) were FACS-purified and sequenced. Following QC, 8483 nuclei (macaque) or 21907 (pig), 5965 (hamster), 7457 (chicken) and 4957 (zebrafish) cells were analyzed using TotalVI (pig) or scVI (macaque, hamster, chicken, zebrafish) (top). KCs were identified using the human-murine KC signature and the signature finder algorithm (Pont et al., 2019) (bottom).

### An evolutionarily-conserved program of KCs

To date, no validated markers of *bona fide* human KCs have been described. Explaining the difficulty to accurately define human KCs, we found monocytes and macrophages formed a single continuum in the human sc/snRNA-seq data, preventing a simple definition of human KCs (Fig. 4B). To further investigate potential human hepatic macrophage heterogeneity, we zoomed in on myeloid cells, identifying 10 clusters (Figs. 4H, S5H,I & Tables S11,12). To define the KCs, we next examined expression of the top 25 murine KC genes by these clusters which identified cluster10 to be the genuine human KCs. Unlike in mice, these were preferentially located in the mid-zone (Fig. S5J,K). Cluster9 also expressed many of these genes but lacked *TIMD4* (Fig. S5H,J), suggesting that these cells may be recently recruited monocyte-derived KCs (moKCs) (Scott et al., 2016). Although *VSIG4* expression alone did not accurately identify KCs in the human liver, VSIG4 protein was found to be the best marker of human KCs in the CITE-seq data highlighting the importance of examining the surface proteome alongside the transcriptome (Figs. 4I, S5I,J). This was validated by flow cytometry and by co-staining human livers for VSIG4 and the KC-specific gene *CD5L,* which also confirmed their mid-zonal localization (Figs. 4J, S5M,N). To assess if KC identity was further conserved in evolution, we profiled macaque, pig, hamster, chicken and zebrafish livers (Fig. 4K). We identified the KCs in an unbiased manner by mapping the conserved human-mouse KC signature onto the datasets (Figs. 4K, S6A-C). We then examined the main features of each KC population identified (Fig. S6D-H & Tables S13-18). A strong overlap in transcriptomes across species was observed likely due to the conserved expression of core KC transcription factors (Fig. S6I). VSIG4 protein expression was also conserved in pig and macaque KCs (Fig. S6J-L). Similarly, we were also able to identify most of the other hepatic cells across species on the basis of conserved genes (Fig. S6M,N). cDC2s were the main exception to this, as specific cDC2 marker genes were not conserved across all species (Fig. S6N).

### LAM location is altered in the steatotic human liver

Alongside KCs, we also identified human CD68^+^VSIG4^−^ macrophages in the liver capsule, in close proximity to central and portal veins as well as at bile ducts (Fig. 5A-C). Similar populations were also observed at the portal and central veins and at the bile ducts in the healthy macaque liver (Fig. S6L). Examination of the scRNA-seq data and comparison with murine signatures identified immature and mature LAMs (Figs. 4H, 5D) enabling the definition of a conserved LAM signature (Fig. S7A,B). Despite a recent study suggesting these cells are specific to fibrotic livers (Ramachandran et al., 2019), we identified LAMs in all patients profiled with scRNA-seq, but there was a trend towards increased proportions of LAMs in the 2 livers with >10% steatosis (Fig. S7C). Consistent with the microscopy and murine bile-duct LAMs, human LAMs were located in portal zones of non-steatotic livers. However, in steatotic human livers, LAMs were primarily located peri-centrally, correlating with the steatosis (Fig. 5E). This change in location of LAMs was also validated using Visium in the mouse following feeding of a western diet (WD) for 36 weeks to induce fatty liver disease (Remmerie et al., 2020). Here, fitting with the presence of steatosis throughout the liver, LAMs were found across portal, peri-portal and mid zones (Fig. S7D). Given that the locations of LAMs in both species correlated with the presence of excess lipid and the heterogeneity in the niche cells between the distinct locations in which LAMs were found (Fig. 5F-H), this suggests that LAM phenotype and abundance may be regulated by lipid exposure rather than by specific local cell-cell interactions conserved at the two locations.

**Figure 5:**
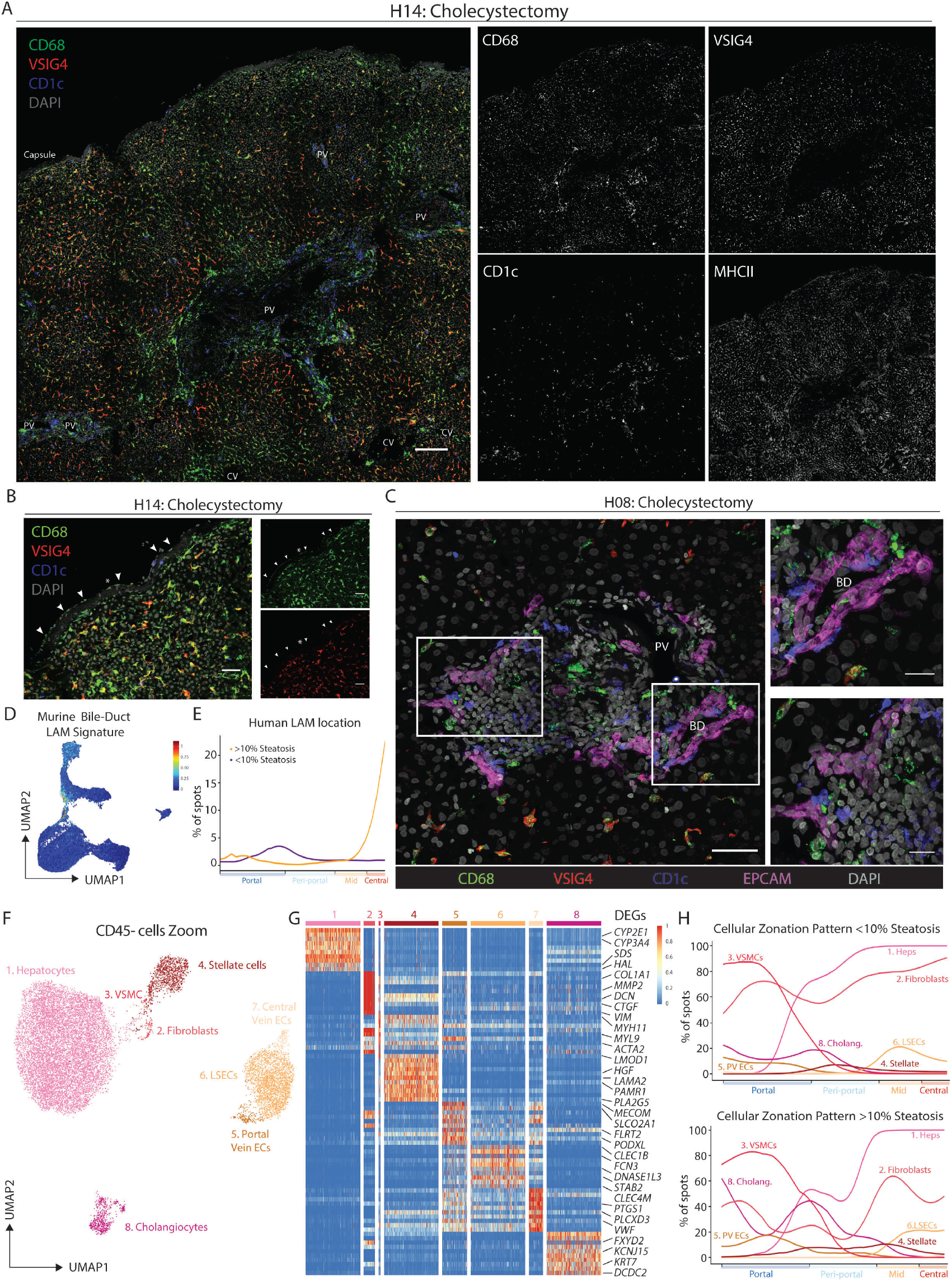
Distinct macrophage populations are also found residing in spatially resolved niches in the healthy human liver. (A,B,C) Expression of indicated markers in human liver (A), capsule (B) or portal tract (C) by confocal microscopy. Scale bar 200μm (A), 20μm (B) and 50μm or 20μm (insets; C). Images are representative of 1-4 patients. PV; portal vein, CV; central vein. Arrowheads indicate capsule macrophages. (D) Human bile-duct LAMs identified using the murine bile-duct LAM gene signature and the signature finder algorithm (Pont et al., 2019). (E) Human LAM signature from scRNA-seq mapped onto the Visium zonation data, healthy (purple), steatotic (orange). (F) UMAP of human CD45^−^ cells (15481 cells/nuclei) isolated from Fig. 4B and re-clustered with scVI. (G) Top DEGs between cell types identified. (H) Indicated cell signatures from sc/snRNA-seq mapped onto the Visium zonation data, healthy (top), steatotic (bottom).

### Differential NicheNet analysis across species reveals a crucial role for the ALK1-BMP9/10 axis in KCs

To assess the roles of conserved cell-cell interactions versus local metabolites, such as lipids, in driving macrophage heterogeneity across species, we performed a differential NicheNet (Browaeys et al., 2019) analysis between the distinct hepatic macrophages and the CD45^−^ cells present in their respective niches focusing on ligands and receptors conserved in both human and mouse (Fig. 6A). In contrast to KCs, this revealed very few specific ligand-receptor pairs for LAMs (Figs. 6A), further hinting that the main signals driving the LAM phenotype may not come from unique cell-cell interactions. Consistent with this, BM-monocytes cultured with acetylated low-density lipoprotein expressed LAM-associated genes (Fig. 6B) demonstrating a dominant role for lipids in inducing the LAM phenotype. Further investigation into the cellular cross-talk forming the blueprint of the KC niche found multiple ligand-receptor pairs to be conserved between human and mouse (Fig. 6C). One of these, an ALK1-BMP9/10 circuit between KCs (ALK1; encoded by *Acvrl1*) and stellate cells (BMP9/10 encoded by *Gdf2/Bmp10* respectively) was found to be conserved in all 7 species and was predicted to control the expression of a number of the conserved KC transcription factors (Figs. 6C,D & S7E,F). To validate this axis, we generated *Fcgr1*-Cre x *Acvrl1*^fl/fl^ mice, eliminating ALK1 from macrophages. This led to an almost complete loss of VSIG4^+^ KCs (Fig. 6E-G, S7G), demonstrating that the evolutionarily-conserved ALK1-BMP9/10 axis is crucial for KC presence in the liver.

**Figure 6:**
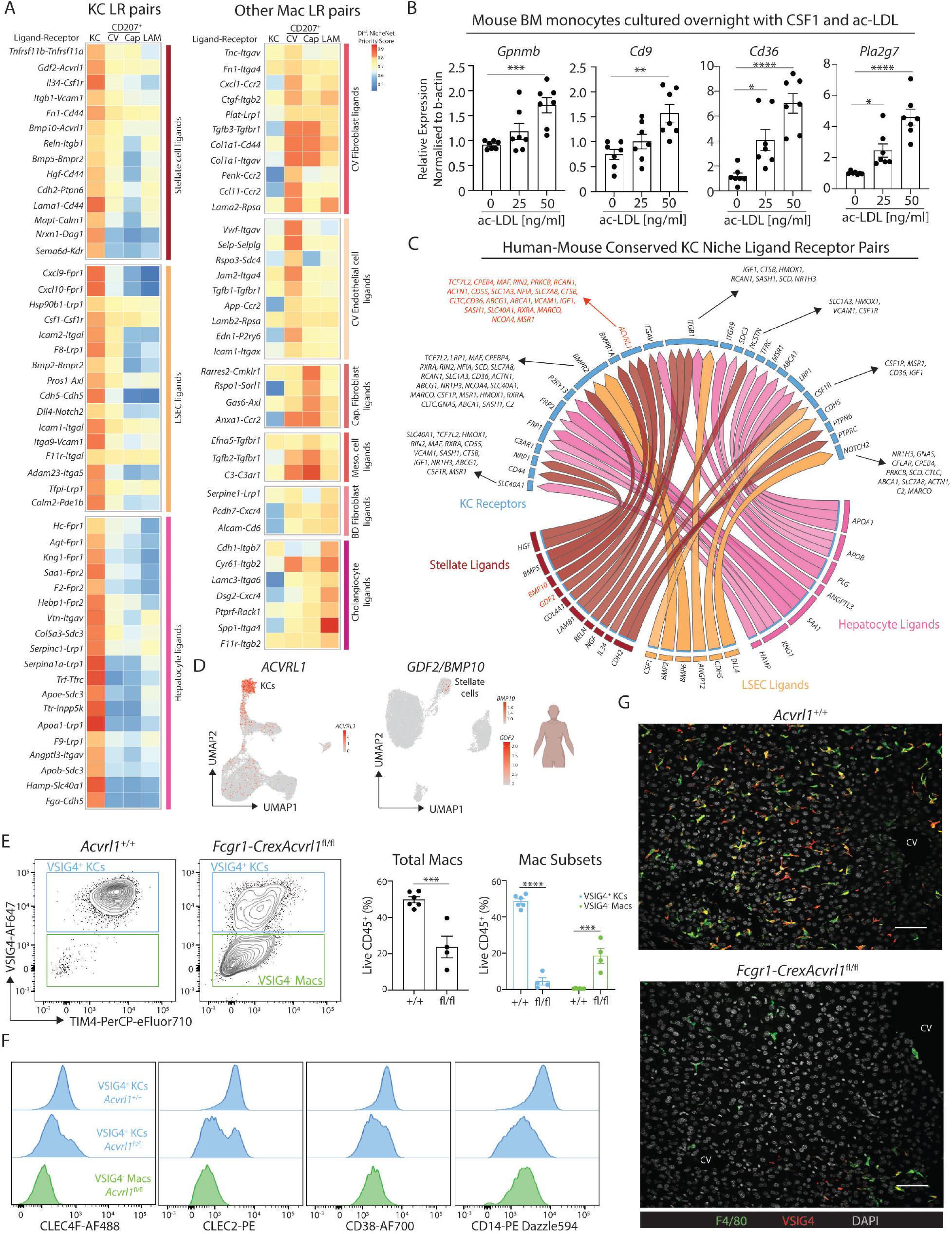
ALK1-BMP9 axis regulates KC presence. (A) Differential NicheNet highlighting prioritized conserved (human-mouse) ligand-receptor (LR) pairs between indicated macrophages and their niche cells. LR pairs are grouped according to the niche cell type with highest ligand expression. (B) Mouse BM monocytes were cultured in the presence of CSF1 and indicated concentrations of ac-LDL, prior to being purified and analyzed for expression of indicated genes by qPCR. Data are pooled from 2 experiments. One-Way ANOVA with Bonferroni post-test compared with 0ng/ml. (C) NicheNet circos plot highlighting conserved ligand-receptor pairs and induced target genes between KCs and indicated niche cells in human and mouse. (D) Feature plots showing expression of ALK1 (*Acvrl1*) in human myeloid cells (left) and *GDF2/BMP10* in CD45^−^ cells (right). (E) Livers were harvested from *Fcgr1*-Crex*Acvrl1*^fl/fl^ mice or *Acvrl1^+/+^* controls and KCs examined (left) and quantified (right) using VSIG4 expression. (F) Expression of indicated KC markers by macrophage populations in *Fcgr1*-Crex*Acvrl1*^fl/fl^ or *Acvrl1^+/+^* control mice. Data are pooled from 2 independent experiments with n=4-5. Student’s t test. (G) Expression of indicated markers in livers of *Fcgr1*-Crex*Acvrl1*^fl/fl^ or *Acvrl1^+/+^* control mice by confocal microscopy. Scale bar 50μm. Images are representative of 2 mice per group. *p<0.05, **p<0.01, ***p<0.001, ****p<0.0001.

## Discussion

To generate a practical cellular atlas of any human tissue and unravel the cell-cell circuits essential for the identities of cells inhabiting that tissue, four key pieces of information are required: (i) an inventory of all cells present, (ii) the location of the different cells within the tissue to identify interactions between neighboring cells, (iii) an alignment between the human and animal models allowing for any predicted cell-cell interactions to be perturbed and (iv) the identification of reliable antibody-based panels for the efficient screening of different patients and/or transgenic animals. Here, by integrating single-cell and spatial transcriptomic and proteomic data, we provide these 4 pieces of information for the liver and uncover evolutionarily-conserved microenvironmental circuits controlling the development of hepatic macrophages.

Highlighting the need to combine different isolation strategies to correctly profile all hepatic cells, we demonstrate that without the combination of scRNA-seq and snRNA-seq, distinct human and murine stromal cell subsets would be lacking from the liver atlas. Unraveling the spatial localization of all hepatic cells, we identify LAMs around the bile-ducts in the healthy mouse, human and macaque liver. However, when steatosis is present, LAMs expand and localize to the steatotic peri-central regions, which is reminiscent of the expansion of LAMs in the livers of obese mice (Remmerie et al., 2020). This spatial information at least partially invalidates the hypothesis that LAM identity is specifically induced by fibrotic stromal cells (Ramachandran et al., 2019) and supports the concept that LAMs are mainly induced by local lipid exposure, as we demonstrate experimentally. Although the precise source of lipid remains to be determined. We also provide an alignment of the liver atlas across seven species. This reveals the conserved transcriptomic program of steady-state KCs and uncovers the spatially-restricted and conserved ligand-receptor pairs between KCs and the cells constituting their niche. Underlining the need to first characterize the healthy tissue before attempting to understand how disease perturbs the cells, we identify the DLL-NOTCH interaction to be an evolutionarily-conserved cross-talk between homeostatic LSECs and KCs and therefore not unique to hepatocellular carcinoma or fibrosis, as recently proposed (Ramachandran et al., 2019; Sharma et al., 2020). Similarly, we find that *FOLR2* expression is not specific to tumor-associated hepatic macrophages (Sharma et al., 2020), but is expressed (mRNA and protein) on KCs in the healthy mouse and human liver. Finally, we apply a proteogenomic pipeline starting from broad oligo-conjugated antibody panels for both single-cell and spatial profiling. This is crucial as transcriptomic profiling does not always correspond with the ability to detect proteins by flow cytometry or microscopy. By screening broadly, we identify the best surface markers for the isolation and localization of hepatic macrophages and their respective niche cells. This allows both the validation of the spatial location at the single-cell level, and the efficient screening of transgenic mouse models for the loss of KCs. Characterization of *Fcgr1*-Crex*Acvrl1*^fl/fl^ mice using our defined panel readily demonstrates the cruciality of the ALK1-BMP9/10 axis in KCs emphasizing that macrophage-fibroblast cross-talk goes much further than the exchange of growth factors (Guilliams et al., 2020; Zhou et al., 2018). Moving forward, applying these relatively cheap antibody panels to large patient cohorts or multiple transgenic mouse models should enable any perturbations disturbing liver homeostasis to be efficiently identified.

## Methods

### Lead Contacts

Further information and requests for resources, data and reagents should be directed to and will be fulfilled by the lead contacts, Charlotte Scott and Martin Guilliams.

### Experimental Model and Subject Details

#### *In Vivo* Animal Studies

##### Mice

WT C57Bl/6J mice (Janvier) were used for this study. Male and Female mice were used for all experiments unless otherwise stated. *Fcgr1-Cre* mice (Scott et al., 2018) were obtained from Prof. Bernard Malissen, CIML, Marseille and crossed with *Acvrl1*^fl/fl^ mice (Park et al., 2008)obtained from Paul Oh, Barrow Neurological Institute, Florida, USA. All mice were used between 6 and 12 weeks of age. All mice maintained at the VIB (Ghent University) under specific pathogen free conditions. All animals were randomly allocated to experimental groups. All experiments were performed in accordance with the ethical committee of the Faculty of Science, UGent and VIB.

##### Pig

Piglets (female, 10 weeks old) were purchased at a local farm and transported to the animal facilities of the Faculty of Veterinary Medicine. The animals were housed in isolation units as blood donors and had access to feed and water ad libitum. At 30 weeks of age, the animals were euthanized by intravenous injection of sodium pentobarbital 20% (60mg/2.5kg) and livers were collected. The animal study was reviewed and approved by the Ethical Committee of the Faculty of Veterinary Medicine (EC2018/55).

##### Chicken

Study animals were clinically healthy Leghorn hens of approximately 58 weeks old collected from a commercial farm. The hens were housed at the Faculty of Veterinary Medicine according to acceptable welfare standards and were observed at least twice daily for health problems. Feed and water was offered *ad libitum*. The chickens were euthanized through intravenous injection (in the wing) with sodium pentobarbital. The EC approval number of this trial was EC2019/015.

##### Macaque

Cynomolgus macaques sourced from China and supplied by Guangzhou Xiangguan Biotech Co., Ltd and confirmed healthy before being assigned to the study. Animal handling, husbandry and euthanasia was performed by WuxiAppTec Co., Ltd., China according to local ethical guidelines (AAALAC accredited 2010). Study animals consists of 4 groups, orally dosed once with a Janssen proprietary immune modulator or vehicle control. Vehicle control animals only were used in this study. Liver tissue samples were snap-frozen immediately after euthanasia. Samples were thawed once before shipping to Ghent for snRNA-seq analysis.

##### Hamster

Female syrian hamsters (Janvier) were housed per one or two in ventilated isolator cages at a temperature of 21°C, humidity of 55% and 12:12 dark/light cycles, with access to food and water *ad libitum* and cage enrichment. All hamsters had SPF status at arrival and manipulations were performed in a laminar flow cabinet. Housing conditions and experimental procedures were approved by the ethical committee of KU Leuven (license P015-2020). Animals were euthanized at 6-8 weeks of age by intraperitoneal injection of 200 mg/mL sodium pentobarbital and livers were collected for analysis.

##### Zebrafish

Zebrafish were maintained under standard conditions, according to FELASA guidelines (Alestrom et al., 2019). All experimental procedures were approved by the ethical committee for animal welfare (CEBEA) from the ULB (Université Libre de Bruxelles) (Protocol #594N). The following transgenic lines at 6 months of age were used: *Tg(mpeg1:EGFP)^gl22^* (Ellett et al., 2011); Tg(*kdrl:Cre*)*^s89^* (Bertrand et al., 2010); *Tg*(*actb2:loxP-STOP-loxP-DsRed^express^*)*^sd5^* (Bertrand et al., 2010) enabling macrophages to be sorted for sequencing as DsRed, GFP double positive cells.

### Patient Studies

Patient studies were run in collaboration with Ghent University Hospital. Liver biopsies (1-2mm^3^) were isolated with informed consent from patients undergoing cholecystectomy or gastric bypass. In addition, liver biopsies were isolated from healthy adjacent tissue removed during liver resection due to colorectal cancer metastasis. In most cases, a second biopsy was also taken to evaluate liver histology. A full overview of all patient samples used in this study can be found in Table S8. All studies were performed in accordance with the ethical committee of the Ghent University Hospital (study numbers: 2015/1334 and 2017/0539).

### Method Details

#### Isolation of Liver Cells

Liver cells were isolated by either *ex vivo* digestion (all species, except zebrafish) or *in vivo* liver perfusion (mice only) and digestion as described previously (Bonnardel et al., 2019; Scott et al., 2016). Briefly, for *ex vivo* digestion, livers were isolated, cut into small pieces and incubated with 1mg/ml Collagenase A and 10U/ml DNAse at 37°C for 20 mins with shaking. For *in vivo* digestion, after retrograde cannulation, livers were perfused for 1-2mins with an EGTA-containing solution, followed by a 5min (6ml/min) perfusion with 0.2mg/ml collagenase A. Livers were then removed, minced and incubated for 20mins with 0.4mg/ml collagenase A and 10U/ml DNase at 37°C. All subsequent procedures were performed at 4°C. Samples were filtered over a 100µm mesh filter and red blood cells were lysed. Samples were again filtered over a 40µm mesh filter. At this point *in vivo* digestion samples only were subjected to two centrifugation steps of 1 min at 50g to isolate hepatocytes. Remaining liver cells (leukocytes, LSECs and HSCs; *in vivo* protocol) and total cells from the *ex vivo* digests were centrifuged at 400g for 5mins before proceeding to antibody staining for flow cytometry.

Dissected livers from 6 months old transgenic zebrafish were triturated and treated with Liberase TM at 33°C for 20 min. Cells were then filtered through 40µm nylon mesh and washed with 2% FBS in PBS by centrifugation. Sytox Red was then added to the samples at a final concentration of 5nM to exclude nonviable cells before proceeding to flow cytometry. DsRed^+^GFP^+^ cells were then FACS-purified.

#### Isolation of Liver Nuclei

Nuclei were isolated from snap frozen liver tissue with a sucrose gradient as previously described (Habib et al., 2016). Briefly, frozen liver tissue is homogenized using Kimble Douncer grinder set in 1ml homogenization buffer with RNAse inhibitors. Homogenised tissue is then is then subjected to density gradient (29% cushion – Optiprep) ultracentrifugation (7700rpm, 4°C, 30 mins). After resuspension, nuclei are stained with DAPI and intact nuclei were FACS-purified from remaining debris.

#### Flow Cytometry and Cell Sorting

Cells were pre-incubated with 2.4G2 antibody (Bioceros) to block Fc receptors and stained with appropriate antibodies at 4°C in the dark for 30-45 minutes. Cell viability was assessed using Fixable Viability dyes (eFluor780 or eFluor506; Thermo Fischer) and cell suspensions were analyzed with a BD FACSymphony or purified using a BD Symphony S6, BD FACSAria II or III. Nuclei were sorted on basis of DAPI positivity and size. Analysis was performed with FlowJo software (BD). Intracellular staining for CD207 was performed by fixing and permeabilizing extracellularly stained cells according to the manufacturer’s instructions using the FoxP3 Fixation/Permeabilization Kit (Thermo Fischer).

#### Confocal microscopy

Confocal staining was performed as described previously (Bonnardel et al., 2019). Immediately after sacrificing mice with CO_2_, inferior vena cava were cannulated and livers were perfused (4 mL/min) with Antigenfix (Diapath) for 5 min at room temperature. After excision, 2-3 mm slices of livers were fixed further by immersion in Antigenfix for 1h at 4°C, washed in PBS, infused overnight in 34% sucrose and frozen in Tissue-Tek OCT compound (Sakura Finetek). 20 µm-thick slices cut on a cryostat (Microm HM 560, Thermo Scientific) were rehydrated in PBS for 5 min, permeabilized with 0,5% saponin and non-specific binding sites were blocked for 30 min with 2% bovine serum albumin, 1% fetal calf serum and 1% donkey or goat serum for 30 minutes. Tissue sections were labeled overnight at 4°C with primary antibodies followed by incubation for 1h at room temperature with secondary antibodies. When two rat antibodies were used on the same section, the directly conjugated rat antibody was incubated for 1h after staining with the unconjugated and anti-rat secondary and after an additional blocking step with 1% rat serum for 30 minutes. Slides were mounted in ProLong Diamond, imaged with a Zeiss LSM780 confocal microscope (Carl Zeiss, Oberkochen, Germany) with spectral detector and using spectral unmixing and analyzed using ImageJ and QuPath software.

#### Confocal Microscopy combined with RNAScope

Experiments were performed using the RNAScope Multiplex Fluorescent V2 Assay kit (ACDBio 323100). Probes targeting intronic regions for *Hs-Cd5l* (ACDBio 850511), Mfa-*Cd5l* (ACDBio 873211), Mm*-Cd5l* (ACDBio 573271), *Mm-Flt3* (ACDBio 487861), *Mm-Xcr1* (ACDBio 562371), *Mm-Mafb* (ACDBio 438531) and *Mm-Mgl2-O1* (ACDBio 822901) were custom-designed and synthesized. They were then labelled with TSA opal 520 (PerkinElmer FP1487001KT), TSA opal 540 (PerkinElmer FP1494001KT), TSA opal 570 (PerkinElmer FP1488001KT), TSA opal 620 (PerkinElmer FP1495001KT) or TSA opal 650 (PerkinElmer FP1496001KT). Tissues were fixed for 16 hours in AntigenFix (Diapath P0016), dehydrated and embedded in OCT as described above. Slices were pre-treated with hydrogen peroxide for 10 min and protease III for 20 min. The recommended Antigen retrieval step was not performed in order to preserve epitope integrity. Probes were hybridized and amplified according to the manufacturer’s instructions. Slides were then stained for protein markers as described above.

#### Visium

Mice were euthanized by means of carbon dioxide (CO_2_) overdose. The liver was excised and consequently trimmed, on ice, to smaller tissue pieces fitting the 10X Visium capture area. Trimmed tissue pieces were embedded in Tissue-Tek® O.C.T.™ Compound (Sakura) and snap frozen in isopentane (Sigma) chilled by liquid nitrogen. Embedded tissue pieces where stored at - 80°C until cryosectioning.

A 10X Visium Spatial Gene expression slide was placed in the cryostat (Cryostar NX70 Thermo Fisher) 30 minutes prior to cutting. 10 µm sections where cut and placed within the capture area. Single 10X Visium Spatial Gene expression slides were stored in an airtight container at −80°C until further processing.

10X Visium cDNA libraries were generated according the manufacturer’s instructions. In short: Tissue sections where fixed in chilled Methanol. A H&E staining was performed to assess tissue morphology and quality. Tissue was lysed and reverse transcription was performed followed by second strand synthesis and cDNA denaturation. cDNA was transferred to a PCR tube and concentration was determined by qPCR. Spatially barcoded, full length cDNA was amplified by PCR. Indexed sequencing libraries where generated via End Repair, A-tailing, adaptor ligation and sample index PCR. Full length cDNA and indexed sequencing libraries were analyzed using the Qubit 4 fluorometer (Thermo Fisher) and Agilent 2100 BioAnalyzer.

#### Visium Highly Multiplexed Protein

Liver slices were prepared as described above for the classical Visium protocol. Slices were dried for 1 min at 37°C and subsequently fixed using 1% paraformaldehyde in PBS. Next, slices were blocked for 30 min (2% BSA, 0.1ug/ul Salmon Sperm, 0.5% Saponin, 1 U/µl protector RNase inhibitor (Roche) in 3X SSC) and incubated with the oligo-conjugated antibody staining mix (2% BSA, 0.1µg/µl Salmon Sperm, 0.5% Saponin, 1 U/µl protector RNase inhibitor, 10uM polyT-blocking oligo (TTTTTTTTTTTTTTTTTTTTTTT*T*T*/3InvdT/), in 3X SSC) for 1h at 4°C. Slides were mounted (90% glycerol, 1 U/µl protector RNase inhibitor) and imaged on Zeiss Axioscan Z1 at 20X magnification. Samples were then processed for a transcriptomic experiment as per manufacturer’s instructions (Visium, 10X Genomics) with modifications to also capture antibody tags. In short, tissue was permeabilized using Tissue Removal Enzyme (Tissue Optimization kit, 10x Genomics) for 9 minutes, as determined by a tissue optimization experiment (10X Genomics, Visium Spatial Tissue Optimization). After reverse transcription, 2 µl of 100 µM FB additive primer (CCTTGGCACCCGAGAATT*C*C*A) per sample was added to the second strand synthesis mix. During cDNA amplification 1 µl of 0,2 µM FB additive primer (CCTTGGCACCCGAGAATT*C*C*A) was added. After cDNA amplification, antibody products and mRNA derived cDNA were separated by 0.6X SPRI select. The purified full-length cDNA fraction was quantified by qRT-PCR using KAPA SYBR FAST-qPCR kit on a PCR amplification and detection instrument. After enzymatic fragmentation indexed sequencing libraries were generated via End Repair, A-Tailing, adaptor ligation and sample index PCR. The supernatant containing antibody product was cleaned up by two rounds of 1.9X SPRI select. Next, 45 µl of the purified antibody fraction was amplified with a 96 deep well reaction module: 95°C for 3 min; cycled 8 times: 95°C for 20 s, 60°C for 30 s, and 72°C for 20 s; 72°C for 5 min; end at 4°C. ADT libraries were purified once more with 1.6X SPRI select. Full length cDNA, indexed cDNA libraries and antibody libraries were analyzed using the Qubit 4 fluorometer (Thermo Fisher) and Agilent 2100 Bioanalyzer. The separation of the cDNA and ADT libraries were performed according to the manufacturer’s instructions (10X genomics).

#### MICS (MACSima™ Imaging Cyclic Staining) technology on the MACSima™ Imaging System by Miltenyi Biotec B.V. & Co. KG

The MACSima™ Imaging System is a fully automated instrument combining liquid handling with widefield microscopy for cyclic immunofluorescence imaging. In brief, staining cycles consisted of the following automated steps: immunofluorescent staining, sample washing, multi-field imaging, and signal erasure (photobleaching or REAlease).

Cryosectioned slices on slides were taken out of the −80°C storage and the appropriate MACSWell™ imaging frame was mounted immediately on the slide. An appropriate volume of ice-cold 4% PFA solution was added (according to the MACSWell™ imaging frames datasheet) and incubated for 10 minutes at room temperature. The slide was washed three times with MACSima Running Buffer. After washing the appropriate initial sample volume of MACSima Running Buffer was added (according to the MACSWell™ imaging frames datasheet). Right before the start of the MACSima ™ instrument a DAPI pre-staining was performed: the MACSima Running Buffer was removed from the sample to be analysed and stained for 10 min with a 1:10 dilution of a DAPI staining solution (volume depends on working volume for the different MACSwell™ formats, see datasheet). The DAPI staining solution was removed and 3 washing steps were performed (MACSima Running Buffer). Finally, the initial sample volume of MACSima Running Buffer was added.

#### Molecular Cartography™

##### Tissue sections

Liver was frozen and sectioned as described above for Visium analysis and liver slices were placed within capture areas on Resolve BioScience slides. Samples were then sent to Resolve BioSciences on dry ice for analysis. Upon arrival, tissue sections were thawed and fixed with 4% v/v Formaldehyde (Sigma-Aldrich F8775) in 1x PBS for 30 min at 4 °C. After fixation, sections were washed twice in 1x PBS for two min, followed by one min washes in 50% Ethanol and 70% Ethanol at room temperature. Fixed samples were used for Molecular Cartography™ (100-plex combinatorial single molecule fluorescence in-situ hybridization) according to the manufacturer’s instructions (protocol 3.0; available for download from Resolve’s website to registered users), starting with the aspiration of ethanol and the addition of buffer BST1 (step 6 and 7 of the tissue priming protocol). Briefly, tissues were primed followed by overnight hybridization of all probes specific for the target genes (see below for probe design details and target list). Samples were washed the next day to remove excess probes and fluorescently tagged in a two-step color development process. Regions of interest were imaged as described below and fluorescent signals removed during decolorization. Color development, imaging and decolorization were repeated for multiple cycles to build a unique combinatorial code for every target gene that was derived from raw images as described below.

##### Probe Design

The probes for 100 genes were designed using Resolve’s proprietary design algorithm. Briefly, the probe-design was performed at the gene-level. For every targeted gene all full-length protein-coding transcript sequences from the ENSEMBL database were used as design targets if the isoform had the GENCODE annotation tag ‘basic’ (Frankish et al., 2018; Yates et al., 2019). To speed up the process, the calculation of computationally expensive parts, especially the off-target searches, the selection of probe sequences was not performed randomly, but limited to sequences with high success rates. To filter highly repetitive regions, the abundance of *k*-mers was obtained from the background transcriptome using *Jellyfish* (Marçais and Kingsford, 2011). Every target sequence was scanned once for all *k*-mers, and those regions with rare *k*-mers were preferred as seeds for full probe design. A probe candidate was generated by extending a seed sequence until a certain target stability was reached. A set of simple rules was applied to discard sequences that were found experimentally to cause problems.

After these fast screens, every kept probe candidate was mapped to the background transcriptome using *ThermonucleotideBLAST* (Gans and Wolinsky, 2008) and probes with stable off-target hits were discarded. Specific probes were then scored based on the number of on-target matches (isoforms), which were weighted by their associated APPRIS level (Rodriguez et al., 2018), favoring principal isoforms over others. A bonus was added if the binding-site was inside the protein-coding region. From the pool of accepted probes, the final set was composed by greedily picking the highest scoring probes.

The following table highlights the gene names and Catalogue numbers for the specific probes designed by Resolve BioSciences.

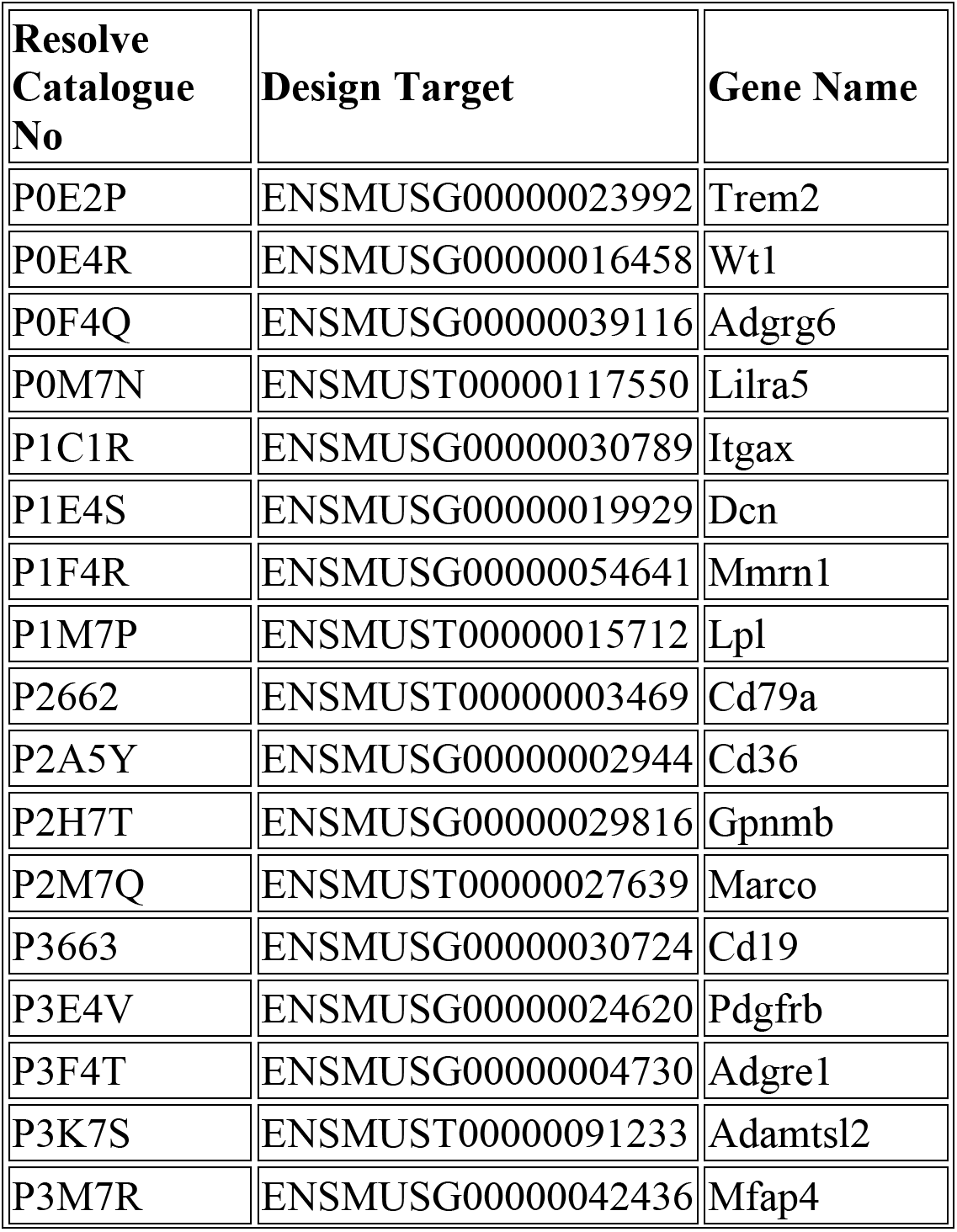
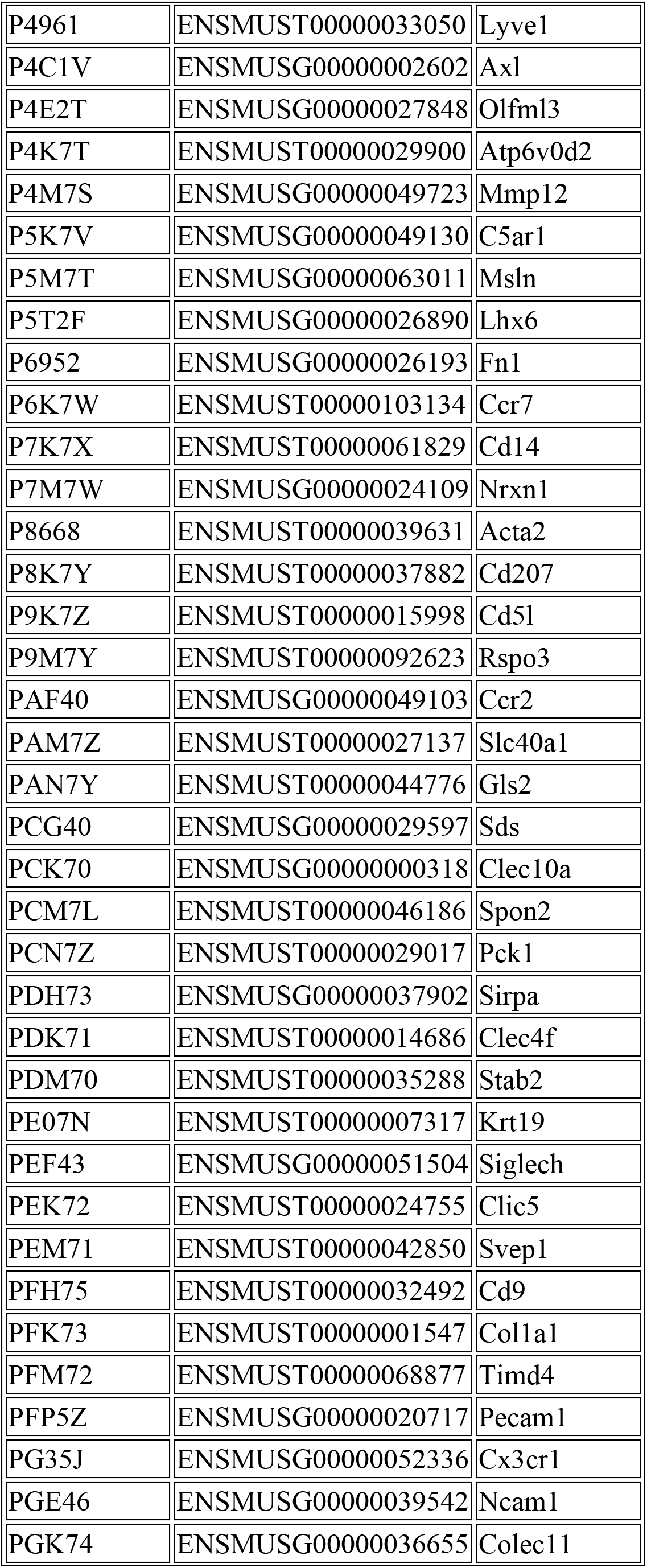
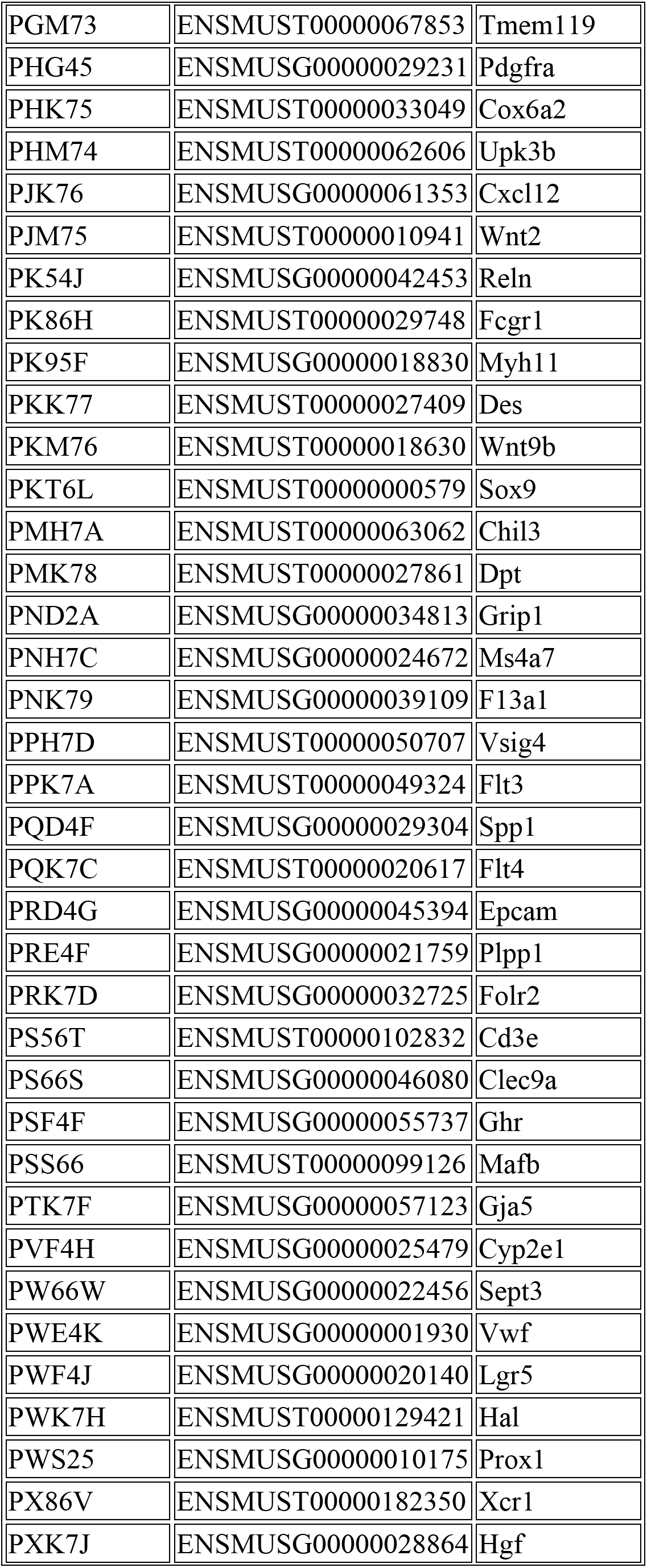
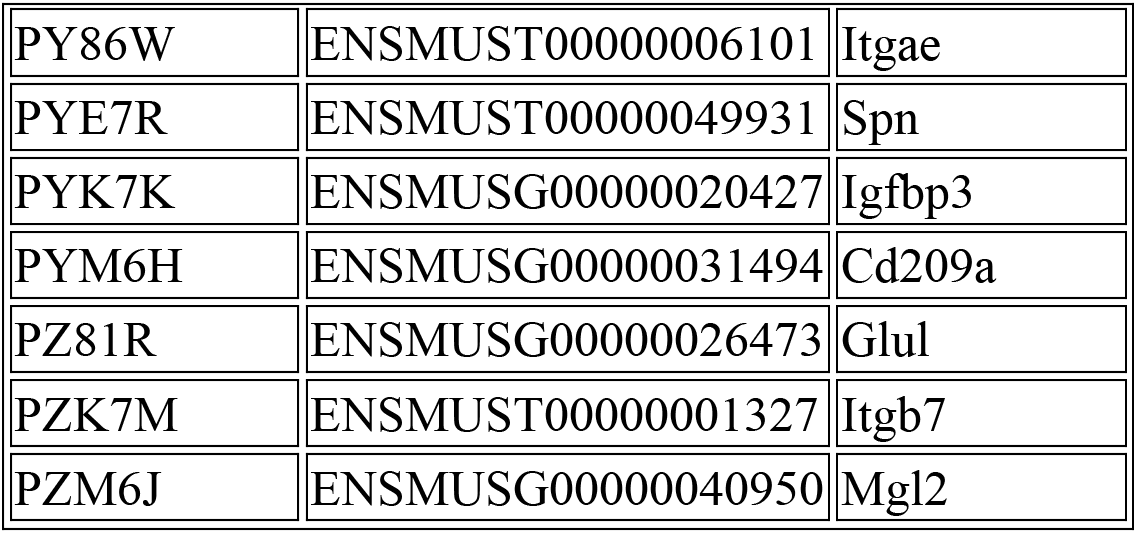

##### Imaging

Samples were imaged on a Zeiss Celldiscoverer 7, using the 50x Plan Apochromat water immersion objective with an NA of 1.2 and the 0.5x magnification changer, resulting in a 25x final magnification. Standard CD7 LED excitation light source, filters, and dichroic mirrors were used together with customized emission filters optimized for detecting specific signals. Excitation time per image was 1000 ms for each channel (DAPI was 20 ms). A z-stack was taken at each region with a distance per z-slice according to the Nyquist-Shannon sampling theorem. The custom CD7 CMOS camera (Zeiss Axiocam Mono 712, 3.45 µm pixel size) was used.

For each region, a z-stack per fluorescent color (two colors) was imaged per imaging round. A total of 8 imaging rounds were done for each position, resulting in 16 z-stacks per region. The completely automated imaging process per round (including water immersion generation and precise relocation of regions to image in all three dimensions) was realized by a custom python script using the scripting API of the Zeiss ZEN software (Open application development).

##### Spot Segmentation

The algorithms for spot segmentation were written in Java and are based on the ImageJ library functionalities. Only the iterative closest point algorithm is written in C++ based on the libpointmatcher library (https://github.com/ethz-asl/libpointmatcher).

##### Preprocessing

As a first step all images were corrected for background fluorescence. A target value for the allowed number of maxima was determined based upon the area of the slice in µm² multiplied by the factor 0.5. This factor was empirically optimized. The brightest maxima per plane were determined, based upon an empirically optimized threshold. The number and location of the respective maxima was stored. This procedure was done for every image slice independently. Maxima that did not have a neighboring maximum in an adjacent slice (called z-group) were excluded. The resulting maxima list was further filtered in an iterative loop by adjusting the allowed thresholds for (Babs-Bback) and (Bperi-Bback) to reach a feature target value (Babs: absolute brightness, Bback: local background, Bperi: background of periphery within 1 pixel). This feature target values were based upon the volume of the 3D-image. Only maxima still in a z-group of at least 2 after filtering were passing the filter step. Each z-group was counted as one hit. The members of the z-groups with the highest absolute brightness were used as features and written to a file. They resemble a 3D-point cloud.

##### Final signal segmentation and decoding

To align the raw data images from different imaging rounds, images had to be corrected. To do so the extracted feature point clouds were used to find the transformation matrices. For this purpose, an iterative closest point cloud algorithm was used to minimize the error between two point-clouds. The point clouds of each round were aligned to the point cloud of round one (reference point cloud). The corresponding point clouds were stored for downstream processes. Based upon the transformation matrices the corresponding images were processed by a rigid transformation using trilinear interpolation.

The aligned images were used to create a profile for each pixel consisting of 16 values (16 images from two color channels in 8 imaging rounds). The pixel profiles were filtered for variance from zero normalized by total brightness of all pixels in the profile. Matched pixel profiles with the highest score were assigned as an ID to the pixel.

Pixels with neighbors having the same ID were grouped. The pixel groups were filtered by group size, number of direct adjacent pixels in group, number of dimensions with size of two pixels. The local 3D-maxima of the groups were determined as potential final transcript locations. Maxima were filtered by number of maxima in the raw data images where a maximum was expected. Remaining maxima were further evaluated by the fit to the corresponding code. The remaining maxima were written to the results file and considered to resemble transcripts of the corresponding gene. The ratio of signals matching to codes used in the experiment and signals matching to codes not used in the experiment were used as estimation for specificity (false positives).

##### Downstream Analysis

Final image analysis was performed in ImageJ using genexyz Polylux tool plugin from Resolve BioSciences to examine specific Molecular Cartography™ signals..

#### RNA Sequencing, CITE-seq and qPCR

##### Sorting and RNA Isolation

40000-160000 cells/nuclei of interest (live, live CD45^+^, live CD45^−^, live myeloid cells from livers of the different species were purified, centrifuged at 400g for 5 mins. When CITE-seq was to be performed, cells were then stained with 2.4G2 antibody to block Fc receptors and CITE-seq antibodies for 20mins at 4°C, before being washed in excess PBS with 2% FCS and 2mM EDTA. Cells were then resuspended in PBS with 0.04%BSA at ∼1000 cells/ml. Cell suspensions (target recovery of 8000-10000 cells) were loaded on a GemCode Single-Cell Instrument (10x Genomics, Pleasanton, CA, USA) to generate single-cell Gel Bead-in-Emulsions (GEMs). Single-cell RNA-Seq libraries were prepared using GemCode Single-Cell 3ʹGel Bead and Library Kit (10x Genomics, V2 and V3 technology) according to the manufacturer’s instructions. Briefly, GEM-RT was performed in a 96-Deep Well Reaction Module: 55°C for 45min, 85°C for 5 min; end at 4°C. After RT, GEMs were broken down and the cDNA was cleaned up with DynaBeads MyOne Silane Beads (Thermo Fisher Scientific, 37002D) and SPRIselect Reagent Kit (SPRI; Beckman Coulter; B23318). cDNA was amplified with 96-Deep Well Reaction Module: 98°C for 3 min; cycled 12 times : 98°C for 15s, 67°C for 20 s, and 72°C for 1 min; 72°C for 1 min; end at 4°C. Amplified cDNA product was cleaned up with SPRIselect Reagent Kit prior to enzymatic fragmentation. Indexed sequencing libraries were generated using the reagents in the GemCode Single-Cell 3ʹ Library Kit with the following intermediates: (1) end repair; (2) A-tailing; (3) adapter ligation; (4) post-ligation SPRIselect cleanup and (5) sample index PCR. Pre-fragmentation and post-sample index PCR samples were analyzed using the Agilent 2100 Bioanalyzer.

##### qPCR

RNA was extracted from 10000 sorted cells (gated using strategies shown) from livers of C57BL/6 mice using a RNeasy Plus micro kit (QIAGEN). Sensifast cDNA synthesis kit (Bioline) was used to transcribe total RNA to cDNA. Real-time RT-PCR using SensiFast SYBR No-Rox kit (Bioline) was performed to determine gene expression, therefore a PCR amplification and detection instrument LightCycler 480 (Roche) was used. Gene expression was normalized to β-actin gene expression. Primers used in the study are detailed below:

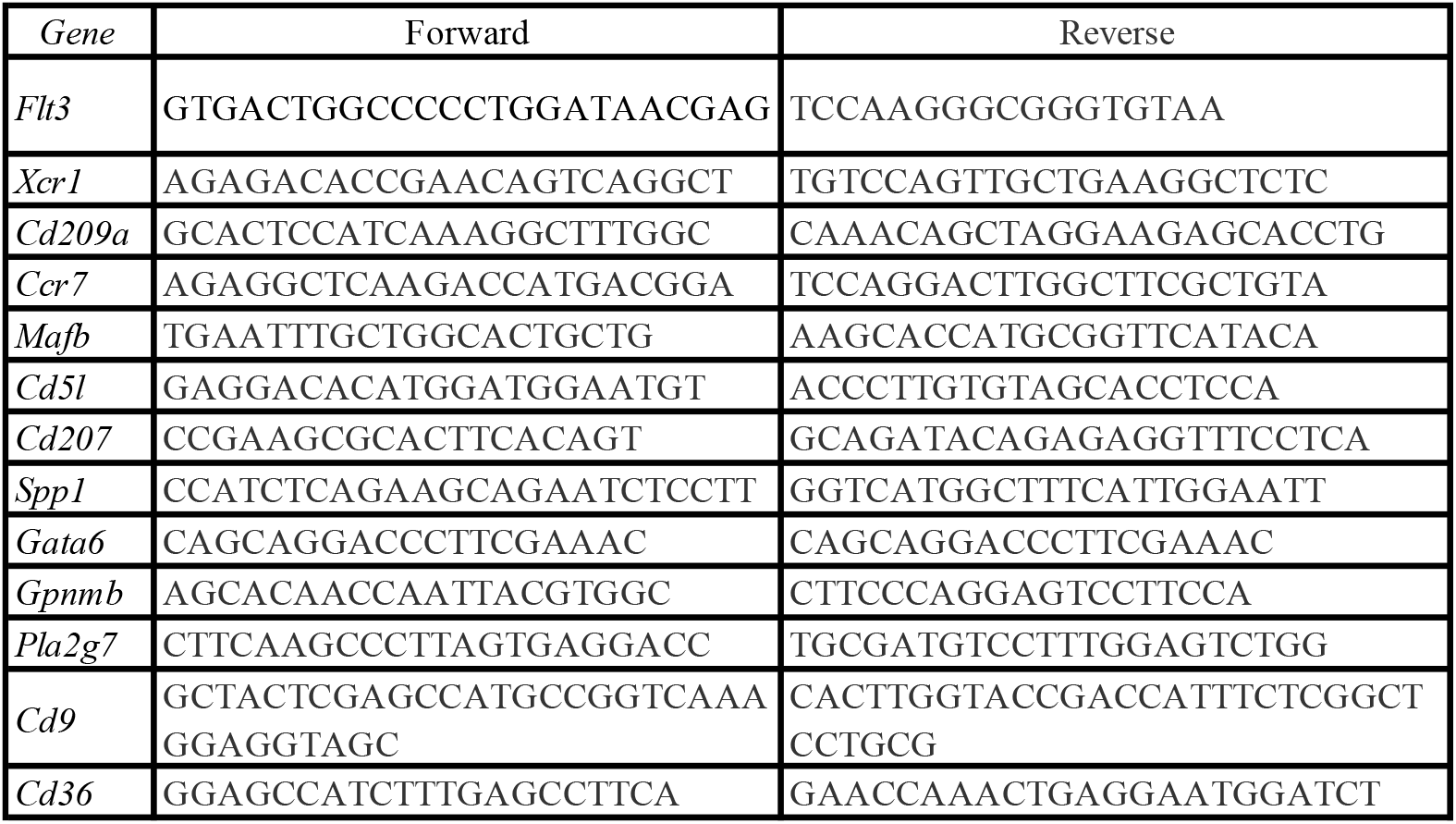

##### RNA Sequencing Analysis

sc/snRNA-seq libraries were loaded on an Illumina HiSeq or Illumina NovaSeq 6000 with sequencing settings recommended by 10X Genomics (26/8/0/98 – 2.1pM loading concentration, ADT and cDNA libraries were pooled in a 25:75 ratio). Visium sequencing libraries were loaded on an Illumina NovaSeq 6000 with sequencing settings recommended by 10X Genomics (28/10/10/75 – 2.1pM loading concentration). Sequencing was performed at the VIB Nucleomics Core (VIB, Leuven). The demultiplexing of the raw data was performed using CellRanger software (10x – version 3.1.0; cellranger mkfastq which wraps Illumina’s bcl2fastq). The reads obtained from the demultiplexing were used as the input for ‘cellranger count’ (CellRanger software), which aligns the reads to the mouse reference genome (mm10) using STAR and collapses to unique molecular identifier (UMI) counts. The result is a large digital expression matrix with cell barcodes as rows and gene identities as columns.

##### Preprocessing Data

To remove ambient RNA, the FastCAR R package (v0.1.0) with a contamination chance cutoff of 0.05 was run on the samples separately before merging them. The UMI cut off was determined individually for the different samples, using the CellRanger web_summary output plot (see GitHub). The Scater R package (v1.14.6) was used for the preprocessing of the data. The workflow to identify the outliers, based on 3 metrics (library size, number of expressed genes and mitochondrial proportion) described by the Marioni lab (Lun et al., 2016) was followed. As a first step cells with a value x median absolute deviation (MADs) higher or lower than the median value for each metric were removed. This value was determined individually for the different datasets (see github). Secondly, the runPCA function (default parameters) of the Scater R package was used to generate a principal component analysis (PCA) plot. The outliers in this PCA plot were identified by the R package mvoutlier. By creating the Seurat object, genes that didn’t have an expression in at least 3 cells were removed. To normalize, scale and detecting the highly variable genes, the R package SCTransform (v0.2.1) was used. If batch correction (on sample level) was needed, the NormalizeData (log2 transformation), FindVariableFeatures and ScaleData functions of the Seurat R package (v3.1.2) were used in combination with the Harmony R package (v1.0). The Seurat pipeline was followed to find the clusters and create the UMAP plots. The number of principal components used for the clustering and the resolution were determined individually for the different datasets (see GitHub). On these initial UMAP plots we did multiple rounds of cleaning by removing proliferating and contaminating (e.g. doublets) cells. For non CITE-seq datasets the count data for the clean cells acquired by the previous steps were further processed with the scVI model (scvi Python package v0.6.7) (Lopez et al., 2018). Datasets including Cite-seq samples were further processed with the TotalVI model (Gayoso et al., 2019). The workflows described on scvi-tools.org were followed to generate new UMAPs, DEGs and DEPs. This information was further processed with the pheatmap R package (v1.0.12) to create heatmaps using the normalized values (denoised genes) calculated in the scVI/TotalVI workflow. The plots showing the expression of certain genes or proteins are created with the ggplot2 R package (v3.2.1) with a quantile cut off of 0.01.

For mouse all the ABs from the whitelist (181 ABs) were loaded into TotalVI, while for the other species only the added ABs were loaded into TotalVI. For the ‘human liver-pool of techniques and patients’ we noticed that the batch correction (between samples) faced difficulties for the hepatocytes and stellate cells as the cells all originated from snRNA-Seq samples, while the other cell types originated from both snRNA-seq and scRNA-seq samples. To overcome this issue we randomly allocated 30% of the hepatocytes to scRNA-seq samples which were not CITE-seq samples. We did the same for 30% of the stellate cells.

Heatmaps were made by scaling the normalized values (denoised values; calculated in the scVI/TotalVI workflow) using the scale_quantile function of the SCORPIUS R package (v1.0.7) and the pheatmap R package (v1.0.12). The plots showing the expression of certain genes or proteins were created based on the normalized values (denoised values) using a quantile cutoff of 0.99 and via either the ggplot2 R package (v3.2.1) or the scanpy.pl.umap function of the Scanpy Python package (v1.5.1).

### Conserved human-mouse KC signature

To find the conserved human and mouse KC markers we started by identifying the human KC markers. We mapped the annotation of the human myeloid UMAP on the human pool of techniques/patients UMAP to identify the real KCs in this last UMAP. The real KCs were identified as the top part of the Macrophage cluster. Using this new annotation we then calculated the DE genes and DE proteins for each cluster. Some genes are listed as marker for multiple clusters, only for the cluster where the gene had the highest score (raw_normalized_mean1/raw_normalized_mean2*lfc_mean), the gene was kept as marker. This way we found 110 potential human KC markers. We then created a heatmap of these 110 genes (using denoised gene values scaled between 0 and 1) and filtered this heatmap by removing the genes where the scaled normalized value was higher than 0.50 in more than 30% of the cells of a certain cell type other than KCs. Except for the Macrophages, we only removed a gene when it had a scaled normalized value higher than 0.50 in more than 70% of the macrophages. After this filtering we ended up with 36 human KC markers. Next we converted these human gene symbols into MGI IDs via the BioMart tool on the HGNC website (https://biomart.genenames.org/martform/#!/default/HGNC?datasets=hgnc_gene_mart). We found a MGI ID for 30 genes. We then converted these MGI IDs into mouse gene symbols via the MGI webtool (http://www.informatics.jax.org/batch/).

To identify the mouse KC markers we similarly mapped the annotation of the mouse myeloid UMAP on the mouse pool of techniques UMAP to identify the real KCs in this last UMAP. The real KCs matched with the Macrophage cluster. Similarly as in human, the DE genes for each cluster was calculated and genes listed as marker for multiple clusters were dealt with in a similar way. This way we found 264 potential mouse KC markers. We then removed the genes that had a score (raw_normalized_mean1/raw_normalized_mean2*lfc_mean) lower than 10 and ended up with 214 genes. We then created a heatmap of these 214 genes (using denoised gene values scaled between 0 and 1) and filtered this heatmap by removing the genes where the scaled normalized value was higher than 0.50 in more than 30% of the cells of a certain cell type other than KCs. After this filtering we ended up with 68 mouse KC markers. Next we converted these mouse gene symbols into MGI IDs via the MGI webtool (http://www.informatics.jax.org/batch/). We then converted these MGI IDs into human gene symbols via the BioMart tool on the HGNC website (https://biomart.genenames.org/martform/#!/default/HGNC?datasets=hgnc_gene_mart) and ended up with 60 genes.

At this point we found 30 human KC markers and 60 mouse KC markers. In a next step, we only kept the human KC markers that we identified as a Highly Variable Gene (HVG) in the mouse pool of techniques UMAP (20 genes) and the mouse KC markers that were identified as HVGs in the human pool of the techniques UMAP (30 genes). We next put these 20 mouse KC markers in SingleCellSignatureExplorer (Pont et al., 2019) to see where these genes are enriched in the mouse pool of techniques UMAP. In order to only get an enrichment in the KCs we decided to only use top 10 mouse KC markers (ordered on score), together with *Slc40a1* and *Hmox1*. We then started to add the top human KC markers as long as we keep the enrichment solely in the KCs. This way we ended up with final list of 15 human-mouse conserved KC markers.

We next converted these KC markers into the monkey, pig, chicken or zebrafish orthologs by looking up the human gene symbol on NCBI (https://www.ncbi.nlm.nih.gov/search/) and checking if there is an ortholog of the species of interest listed under the ‘Ortholog’ tab. The found orthologs were then used as input for the SingleCellSignatureExplorer tool.

### Conversion of the CITE-seq data into a flow cytometry file

The protein normalized values (denoised values; calculated in the TotalVI workflow) were converted into an FCS file using the write.FCS function of the flowCore R package (v1.50.0).

### Preprocessing Visium Data

We first removed per sample all spots that were clear outliers compared to the location of the tissue. Each sample was then normalized individually using the SCTransform function of the Seurat R package (v3.2.3) with default parameters. All samples were then merged with the merge function of the Seurat R package (v3.2.3) with default parameters. Next, we determined the HVGs, created a PCA plot, performed clustering and created an UMAP plot as described in the spatial workflow available on the Seurat website (https://satijalab.org/seurat/articles/spatial_vignette.html). Clusters which showed high mitochondrial gene expression were removed. Spots located at the darker parts of the tissue were also removed as these parts are considered to be dead tissue or of bad quality.

### Modelling of Visium data

#### Probabilistic graphical modelling

For modelling the cell type composition and zonation, spatial CITE-seq and transcriptomics data were analyzed using probabilistic graphical models, similar to what is used in tools such as cell2location and scVI. In brief, transcriptomics data was modelled as a NegativeBinomial distribution, parameterized with a mean *μ* and dispersion *θ*, the latter optimized as a free parameter for each gene. Visium Highly Multiplexed Protein data was modelled as a mixture of NegativeBinomials, with a *μ*_background_ and *μ*_foreground_ and a shared dispersion *θ*. The actual foreground/background signal within a modality was modelled as a *ρ* that depends on the latent space, and which is multiplied with the empirical library size to get *μ*. For Visium Highly Multiplexed Protein, *ρ_background_* was modelled as a latent variable specific for each gene. *ρ_foreground_* for Visium Highly Multiplexed Protein and *ρ* for RNA-seq were modelled as deterministic functions depending on the use case as described in the following paragraphs. The posterior of the probabilistic graphical model was inferred using black-box variational inference (Ranganath et al., 2013), in which the variational distribution was specified as a diagonal Normal distribution, transformed into the correct domain using transforms 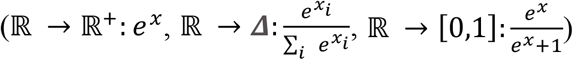. Free parameters within this model were optimized using gradient descent, with the ELBO as loss function and Adam as optimizer as implemented in Pytorch (Paszke et al., 2019) (pytorch.org). We used a learning rate of 0.01 for variational parameters, and 0.001 for parameters of the amortization functions.

### Reference for deconvolution

To calculate the average expression of each gene within a cell type, we used a linear model in which both *ρ* and *θ* were modelled as a latent variable specific for each gene and cell type. The *ρ* for nuclei were multiplied with a gene-specific correction factor (optimized as a latent variable) that corrected for differences between scRNA-seq and snRNA-seq. Given that spatial transcriptomics data sequences the whole cell, the uncorrected *ρ* values were used for spatial deconvolution.

### Deconvolution

To infer the proportions of each cell type within a spot, we used a model in which the gene expression is modelled as a linear combination of cell type proportions and average expression in each cell type:

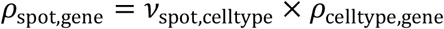

For *ρ*_celltype,gene_ we adapted the values from the reference, but included

- A capture bias per gene, which corrects for technical and biological differences between spatial and sc/sn-RNA-seq. The capture bias was modelled as a latent variable with prior *Normal*(0, 1)
- A red blood cell cell type, which was not included in the reference dataset but nonetheless had a dominant presence in the spatial data. The *ρ* of this cell type was set to zero for all genes except *Hbb-bt*, *Hbb-bs*, *Hba-a1*, *Hba-a2* for mouse and *HBB*, *HBA1*, *HBA2* for human, which were modelled as free parameters.
- Similarly, the expression of complement factors (*C3, C2*, *C4B/C4b*) within hepatocytes was modelled as free parameters.

A background signal shared for all spots was also modelled as follows:

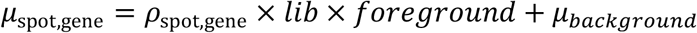

With *foreground* ∈ [0,1] a latent variable specific to each spot and *μ_background_*∈ ℝ a latent variable specific to each gene.

A likelihood ratio test was used to assess whether a cell type was significantly present in a spot. Specifically, if *x* is the gene expression of all genes at a particular spot, we used Monte Carlo samples from the posterior to estimate:

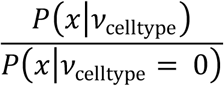

A cell type was deemed significantly present if the log-likelihood was higher than 10.

### Zonation

The zonation of spots was modelled as a univariate latent variable *z* ∼ *Uniform*(0, 1) specific to each spot. This latent variable influenced the gene expression *ρ* using a spline function by using a gaussian basis function (*σ* = 0.05) with 10 knots at uniform fixed positions. The coefficients of this spline were modelled as a latent variable specific for each gene, with prior a Gaussian random walk distribution, and the step ∼ *Normal*(0, **σ*_gene_* ). **σ*_gene_* was determined empirically as 2 times the standard deviation of the log1p transformed expression values in the whole dataset. The variational parameters of the zonation *μ_z_* and *σ_z_* were not optimized directly but were estimated using an amortization function. This amortization function used the count matrix as input, and estimated the variational parameters using the following layers: Linear (with 100 output dimensions), BatchNorm, ReLU, Linear (again with 100 output dimensions), ReLU, and a final Linear layer. This amortization function was used to transfer the zonation onto a different dataset, i.e., 1) to transfer the zonation trained on mouse spatial transcriptomics onto mouse Visium highly multiplexed protein and 2) to transfer the zonation trained on human low steatosis (<10%) onto human high steatosis (>30%).

### Differential abundance along zonation

To determine the differential abundance of a cell type across zonation, the significant presence of a cell type within a spot ∈ {0, 1} was modelled using a spline function with the zonation of a cell type as input. The coefficients of this spline function were modelled as a latent variable with the step size ∼ *Normal*(0, 1). To determine differences in abundance between patients with high and low steatosis, we first modelled the zonation on human data on patients with steatosis < 10%. Potential interaction effects between zonation and steatosis status were then modelled using a spline function as before, but with a separate set of coefficients for both high and low steatosis. A likelihood ratio test was then used to determine whether this interaction was present significantly, by comparing the likelihood of this model with a model with shared coefficients.

#### Differential NicheNet

To analyze cell-cell communication in the hepatic macrophage niches, we applied Differential NicheNet, which is an extension of the default NicheNet pipeline to compare cell-cell interactions between different niches and better predict niche-specific ligand-receptor (L-R) pairs. It uses a flexible prioritization scheme that allows ranking L-R pairs according to several properties, such as niche- and region-specific expression of the L-R pair, ligand activity, and level of database curation. This in contrast to the default NicheNet pipeline which prioritizes expressed L-R pairs solely based on ligand activity predictions. All analyses were conducted according to the Differential NicheNet tutorial (https://github.com/saeyslab/nichenetr/blob/master/vignettes/differential_nichenet.md). As input to the Differential NicheNet pipeline, we used the data after normalization via SCTransform and integration of scRNA-seq and snRNA-seq according to the Seurat procedure for integration (Stuart et al., 2019).

For the mouse analyses, Differential NicheNet was first performed for the following 3 niche comparisons: 1) KCs versus central vein macrophages; 2) KCs versus capsule macrophages; 3) KCs versus LAMs. Following sender cell types were considered for these niches: KC niche: periportal hepatocytes, periportal LSECs, and periportal stellate cells; Central vein macrophage niche: central vein ECs and central vein fibroblasts; Capsule macrophage niche: mesothelial cells and capsule fibroblasts; LAM niche: cholangiocytes and bile duct fibroblasts.

Because of the preferentially periportal localization of KCs in the mouse liver, we also included a ‘region specificity’ factor in the Differential NicheNet prioritization framework. This was done to increase the ranking of ligands that are more strongly expressed in periportal than pericentral niche cells. Periportal sender cells were determined after subclustering based on the following markers: *Hal* and *Sds* for hepatocytes; *Mecom*, *Msr1*, and *Efnb2* for LSECs; *Ngfr*, *Igfbp3*, and *Dach1* for stellate cells.

In the heatmap (Fig. 6A), we show the prioritization scores of the top 40 ligands (and their highest scoring receptor) in the KC niche (score averaged over the 3 analyses), and of all the non-KC niche L-R pairs with a prioritization score ≥ the score of the lowest scoring KC L-R pair of this top 40. For each L-R pair/niche combination, we only displayed the score of the sender cell with the highest score (e.g. for the *Csf1*-*Csf1r* interaction in the KC niche, the score is shown for the LSEC-KC interaction because that score was higher than for Stellate–KC and Hepatocyte–KC; in the LAM niche, the score of *Csf1*-*Csf1r* is shown for the bile duct fibroblast – LAM interaction and not for the cholangiocyte–LAM interaction, etc.).

Because of the strong concordance between the top-ranked L-R pairs in these 3 non-KC macrophage niches, it was decided to also conduct a subsequent analysis in which the KC niche is compared against all non-KC hepatic macrophage niches combined. For this final ‘KC versus all non-KC macrophage analysis’, KCs were compared to central vein macrophages, capsule macrophages, and LAMs together, with the same sender cell types as described here above (but now analyzed together).

For the human analyses, Differential NicheNet was performed to compare the KC niche with the non-KC macrophage niches (similarly as the final analysis in mouse). For the KC niche, all hepatocytes, LSECs, and stellate cells were selected as sender cells; and KCs as receiver cells. For the non-KC macrophage niche, cholangiocytes, fibroblasts, and central vein ECs were considered as the sender cells; Mat. LAMs, Imm. LAMs, and Mac1s as the receiver cells (Fig 4H).

To find KC-niche-specific L-R pairs that are conserved across mouse and human, the individual mouse and human prioritization scores were averaged to form a ‘conservation score’. The 40 ligands (and maximally 3 of their highest scoring receptors) with the highest conservation score were selected for further analysis (note: the L-R pair should be expressed by the same sender-receiver pair in both species). In the circos plot (Fig. 6C) (Gu et al., 2014), only a subset of these top L-R pairs is shown to keep the figure clearly interpretable. Following ligands were not shown: *ITGA9*, *SEMA6D*, *JAM3*, *ITGB1* (stellate cells); *ITGA9*, *F8*, *CD274*, *HSP90B1* (LSECs); *C5*, *F9*, *F2*, *FGA*, *TF*, *TTR*, *COL18A1*, *COL5A3*, *SERPINA1*, *SERPINC1* (hepatocytes). The depicted target genes are KC-specific in both mouse and human, and a top-predicted target according to the NicheNet ligand-target regulatory potential scores. *NR1H3* was manually added as a *NOTCH2* target based on recent studies (Bonnardel et al., 2019).

#### Isolation and culture of BM monocytes with acetylated -LDL

BM was isolated from the tibia and femur of mice by centrifugation. Red blood cells were lysed and single cell suspensions were stained with antibodies for flow cytometry. BM monocytes were sorted as live CD45+ CD11b+ Ly6G- Ly6C+ CD115+ cells using a BD FACSAria III. Monocytes were resuspended in DMEM/F12 media supplemented with 10% FCS, 30ng/ml CSF1, 2mM Glutamine and 100U/ml penicillin and streptomycin. 150,000 monocytes were seeded in each well of an adherent 24-well plate pre-coated with bovine collagen type I and cultured overnight (37C, 5% CO2). The following day 0, 25 or 50ng/ml of ac-LDL was added. 14 hours later cells were harvested and live F4/80+ cells were FACS-purified in RLT plus buffer containing 1% β-mercaptoethanol. RNA isolation, cDNA synthesis and qPCR were performed as described above.

### Quantification and Statistical Analysis

In all experiments, data are presented as mean ±SEM and/or individual data points are presented unless stated otherwise. Statistical tests were selected based on appropriate assumptions with respect to data distribution and variance characteristics. Details of the precise test used for each analysis can be found in the figure legends. Statistical significance was defined as p<0.05. Sample sizes were chosen according to standard guidelines. Number of animals/patients is indicated as ‘‘n’’. The investigators were not blinded to the group allocation, unless otherwise stated.

## Acknowledgments

We thank all patients and their families for participating in this study. We also thank Janssen for providing the Macaque samples, the IRC-VIB Flow and Bioimaging core facilities for assistance, VIB Tech Watch and the VIB Single-cell accelerator program for their help benchmarking technologies including CITE-Seq, Molecular Cartography and MICS, and 10X Genomics for their help setting up the Visium highly-multiplexed protein analysis. Finally, we thank the VIB-UGent animal house staff. BioRender was used to generate some figures.

## Funding

Chan Zuckerberg Initiative; Liver Seed Atlas Grant (MG, CLS)

FWO SBO; iPSC LiMics (CLS, MG, YS)

ERC Consolidator Grant; KupfferCellNiche, 725924 (MG)

GOA; BOF18-GOA-024 (MG, YS)

ERC Starting Grant; MyeFattyLiver, 851908 (CLS)

FWO Project Grants; 3G000519, (CLS, MG)

FWO PhD Fellowship; 1129521N (BH), 1181318N (RB)

MSCA IF Fellowship; MACtivate 101027317 (CZ)

## Author contributions

Conceptualization: CLS, MG

Methodology: CLS, MG, BH, JB, LM, TT, RB, WS, AB, AG, SL

Investigation: CLS, MG, BH, JB, BVDB, AB, FDP, CZ, BV, TV, LM, TT, RB, AH, AV, FB, YV, BD, EC, GF, VW, AW, SK, JN, KD, PG, SC

Visualization: CLS, MG, JB, BH, LM, TT, RB, WS

Funding acquisition: CLS, MG

Supervision: CLS, MG, HVV, LD, WS, BD, YS

Writing: CLS, MG

## Supplementary Data

Figures S1-S7

Tables S1-S19

**Fig. S1:**
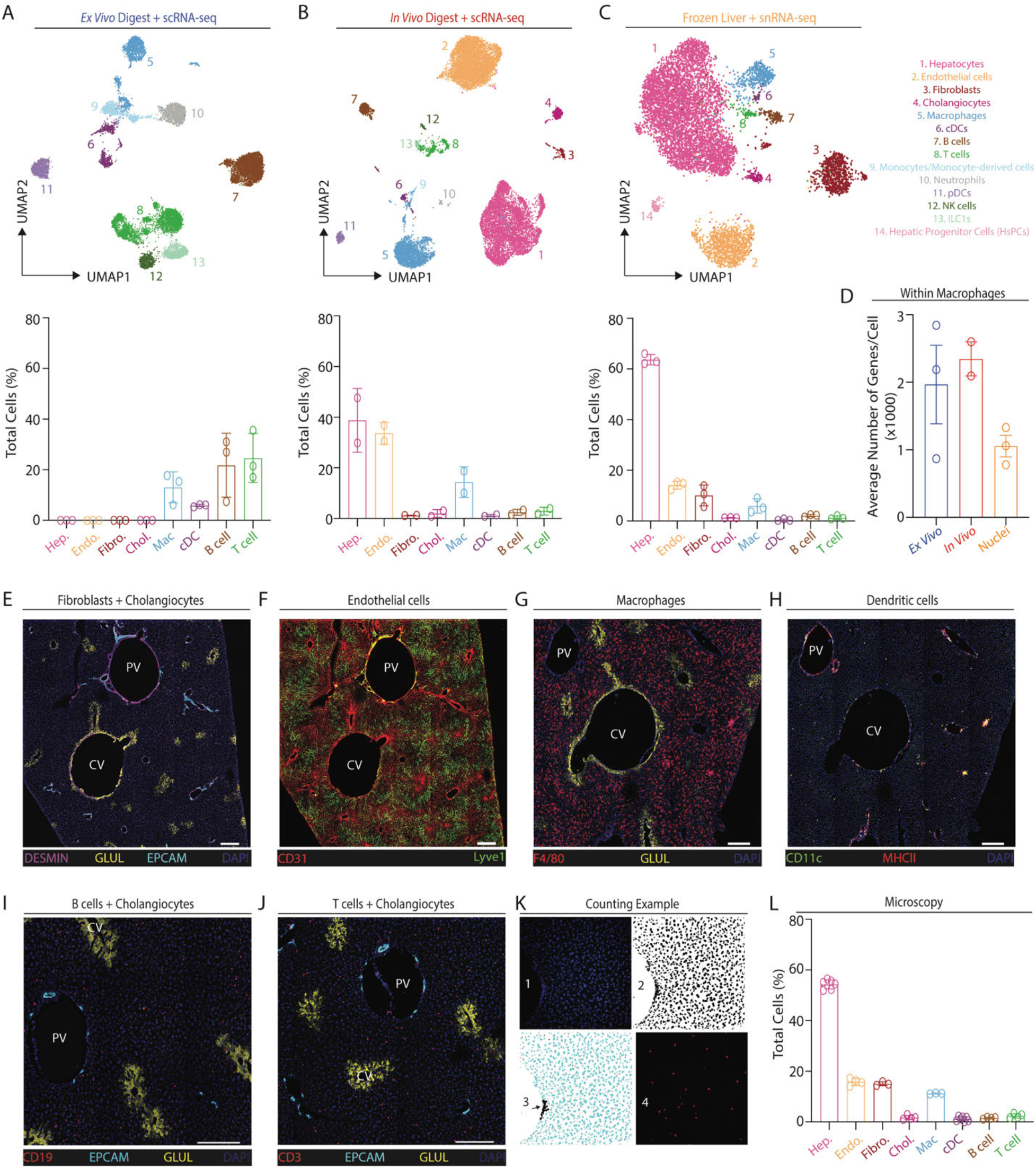
Cell types identified in transcriptomic studies depend upon cell/nuclei isolation technique used. Related to Fig. 1. Cells were isolated from livers of healthy C57B/l6 mice by either *ex vivo* (3 mice) or *in vivo* (2 mice) enzymatic digestion. Alternatively, livers were snap frozen and nuclei subsequently isolated following tissue homogenization by a sucrose gradient (3 mice). Live cells/intact nuclei were identified and purified using flow cytometry. For the cells, either live CD45^+^, live CD45^−^ or live hepatocytes or were sorted. 1 *ex vivo* digested sample and 1 *in vivo* digested sample were also stained with a panel of 107 (*ex vivo* cells) or 161 (*in vivo* cells) oligo-conjugated antibodies for CITE-seq analysis. FACS-purified cells/nuclei were loaded onto the 10X Chromium platform and scRNA-seq, CITE-seq or snRNA-seq performed. Following clean up and QC, cells from the same mice were pooled together in the same ratios (CD45^+^:CD45^−^:Heps) as found in the tissue as a whole before sorting, different mice were then pooled together and the data were analyzed using scVI. (A-C) UMAPs showing annotations of cell-types and proportions of each cell type as a % of total cells in the UMAP isolated using (A) *ex vivo* digestion; 13144 cells, (B) *in vivo* digestion; 19428 cells and (C) nuclei; 8583 nuclei. (D) Average number of genes/cell in the annotated macrophage population following each isolation method. (E-J) Confocal microscopy images to determine true abundance of (E) fibroblasts and cholangiocytes (F) endothelial cells, (G) macrophages, (H) dendritic cells, (I) B cells and (J) T cells *in vivo*. Scale bar 200μm. (K) The percentage of each population was calculated based on the percentage of a given population divided by the total number of nuclei. A threshold was applied to the DAPI channel (picture 1) in ImageJ (picture 2) and nuclei were automatically counted based on the ImageJ ‘analyze particles’ plugin (size (micron\2 = 10-1000; circularity = 0-1; picture 3). Due to the density of some liver zones, some nuclei were not automatically counted (arrow, picture 3). Those were then manually counted and added to the total number of nuclei. For the populations of interest, cells were counted manually based on specific markers (for example, CD3 for T cells, picture 4). Counting was performed blinded prior to analysis of the sequencing results. (L) Proportion of indicated cell types as a % of total cells identified in confocal microscopy images. Data are from 3-7 images per cell type taken from 2-4 mice.

**Fig. S2:**
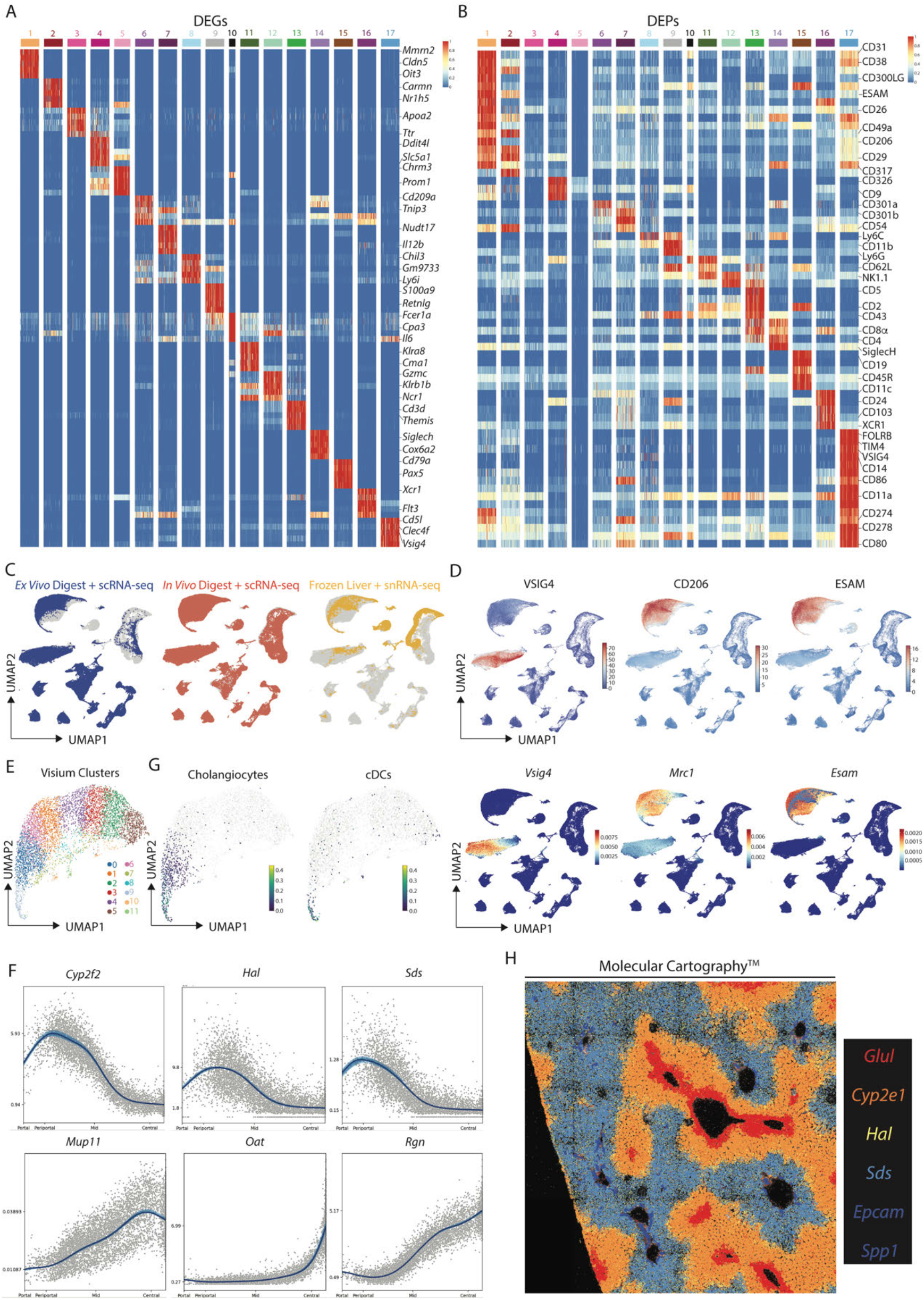
Combination of CITE-seq, scRNA-seq, snRNA-seq and spatial analyses enables generation of a mouse liver atlas. Related to Figure 1. (A,B) Top DEGs (A) and DEPs (B) for cell types identified in Fig. 1B. (C) Distinct profiles of cells or nuclei within the UMAP depending on isolation protocols; 71162 cells from *ex vivo* digestions, 96066 cells from *in vivo* digestions and 18666 nuclei. (D) Expression of VSIG4, CD206 and ESAM (protein, top) and *Vsig4*, *Mrc1* and *Esam* (mRNA, bottom). (E) UMAP showing clusters generated from Visium analysis of liver tissue (4 samples) and liver capsule (1 sample). (F) Top unbiased genes defining zonation trajectory from portal to central vein in Visium. (G) Identification of Cholangiocyte (left) and cDC (right) signatures on zonated Visium spots. (H) Molecular Cartography showing expression of indicated zonated hepatocyte mRNAs in liver tissue. Data are representative of 2 mice.

**Fig. S3:**
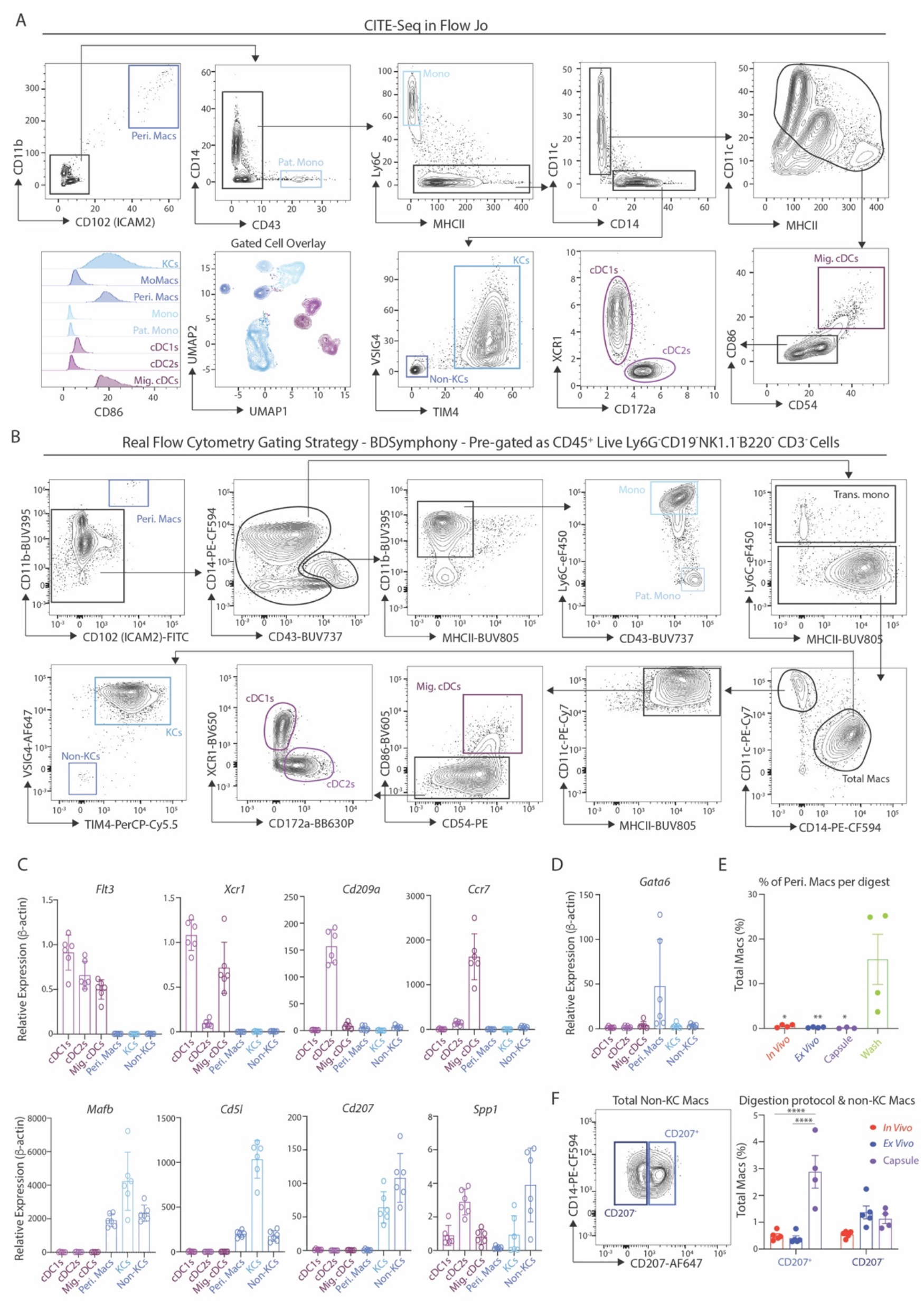
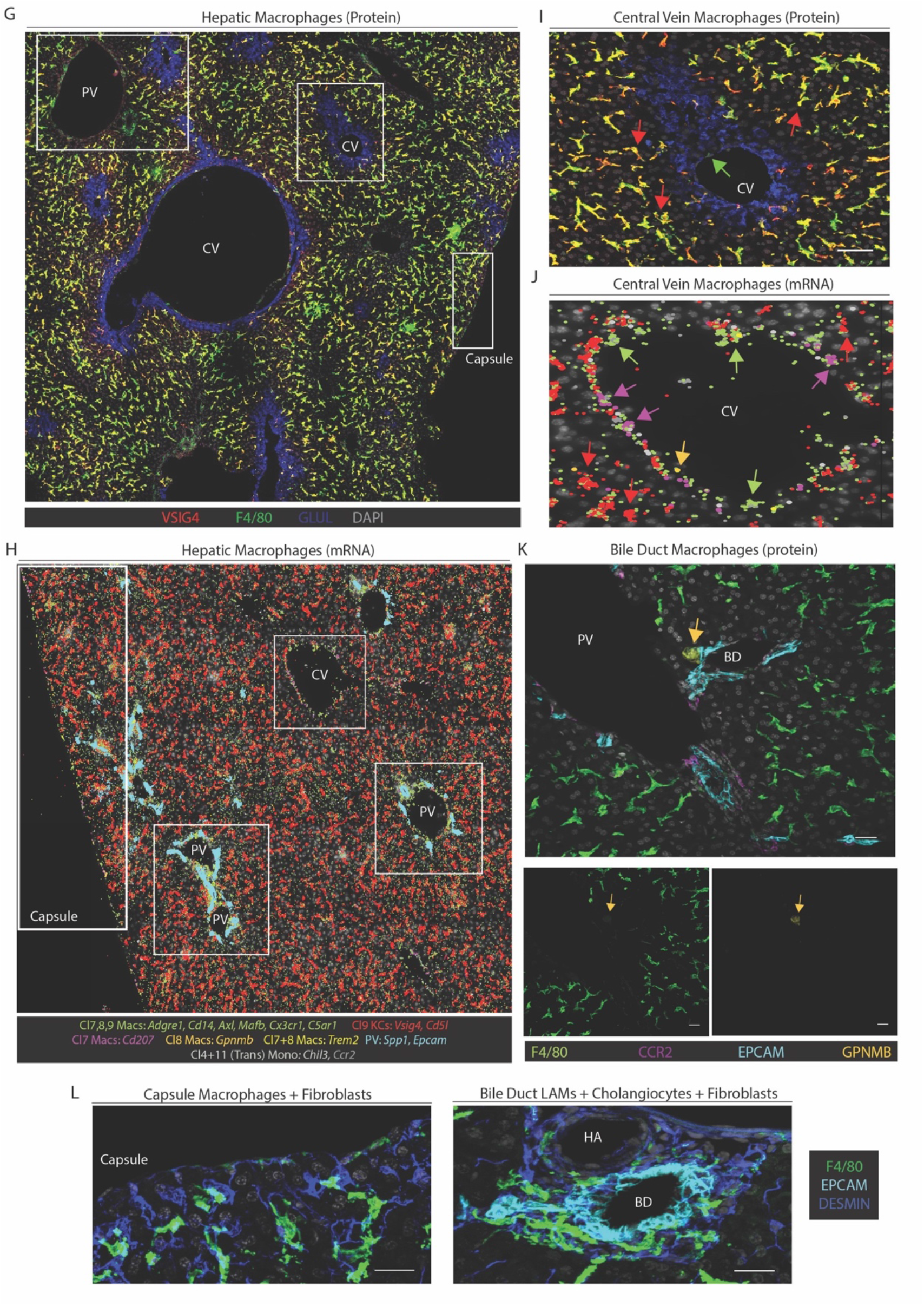
Validated flow cytometry gating strategy for murine myeloid cells. Related to Figure 2. (A) CITE-seq data from the murine myeloid cells in Fig. 2A were exported as an FCS file and an *in-silico* gating strategy identified in FlowJo software. (B) Application of the *in-silico* gating strategy with a 21-colour flow cytometry panel. Myeloid cells were pre-gated as live CD45^+^Lineage-cells (Ly6G^−^CD19^−^NK1.1^−^B220^−^CD3^−^). Data are representative of 3 experiments with 3-6 mice per experiment. (C) cDC1s, cDC2s, Migratory cDCs (Mig. cDCs), Peritoneal macrophages (Peri. Macs), KCs and non-KC macrophages (Non-KCs) were FACS-purified using gating strategy in B, mRNA was isolated and qPCR performed to examine expression of indicated genes defining each population to validate their identity. Data are representative of 2 experiments with n=3-6. (D) Putative peritoneal macrophages were FACS-purified using gating strategy in (B) and expression of *Gata6* was examined by qPCR compared with other hepatic myeloid populations. Data are from a single experiment with n=6. (E) Peritoneal macrophages as a % of total macrophages recovered from the liver using different digestion techniques (*in vivo*, e*x vivo* or capsule) or in supernatants in which livers were washed following removal from the mouse but prior to digestion (wash). Data are from a single experiment with n=4. *p<0.05, **p<0.01 One-way ANOVA with Bonferroni post-test compared with wash data. (F) Expression of CD14 and CD207 within the non-KC macrophage population from B (left) and % of CD207^+^ and CD207^−^ populations amongst total macrophages in livers digested using the *ex vivo* or *in vivo* protocols or in dissected and digested liver capsule (right). Data are representative of two experiments with n=4-5 mice per experiment. ****p<0.0001 mixed effects analysis with Tukeys multiple comparison test. (G) Expression of VSIG4, F4/80, GLUL and DAPI by confocal microscopy. Insets represent zones featured in Fig. 2E,G and Fig. S3I. (H) Molecular Cartography of indicated genes and cell types. Insets represent zones featured in Fig. 2F,H and Fig. S3J. (I) Expression of VSIG4, F4/80, GLUL and DAPI by confocal microscopy at the central vein. Scale bar 50μm. (J) Molecular Cartography of indicated genes and cell types at central vein. (K) Expression of F4/80, EPCAM, CCR2, GPNMB and DAPI by confocal microscopy at a portal vein (top) or F4/80 or GPNMB alone (bottom). Scale bar 25μm. (L) Expression of DESMIN and F4/80 at the liver capsule and underlying parenchyma (left) or EPCAM, DESMIN and F4/80 at the bile duct by confocal microscopy. PV; portal vein, CV; central vein, HA; hepatic artery, BD; bile duct. Arrows indicate specific cell types, where color corresponds to cell type/markers. All images are representative of 2-6 mice.

**Fig. S4:**
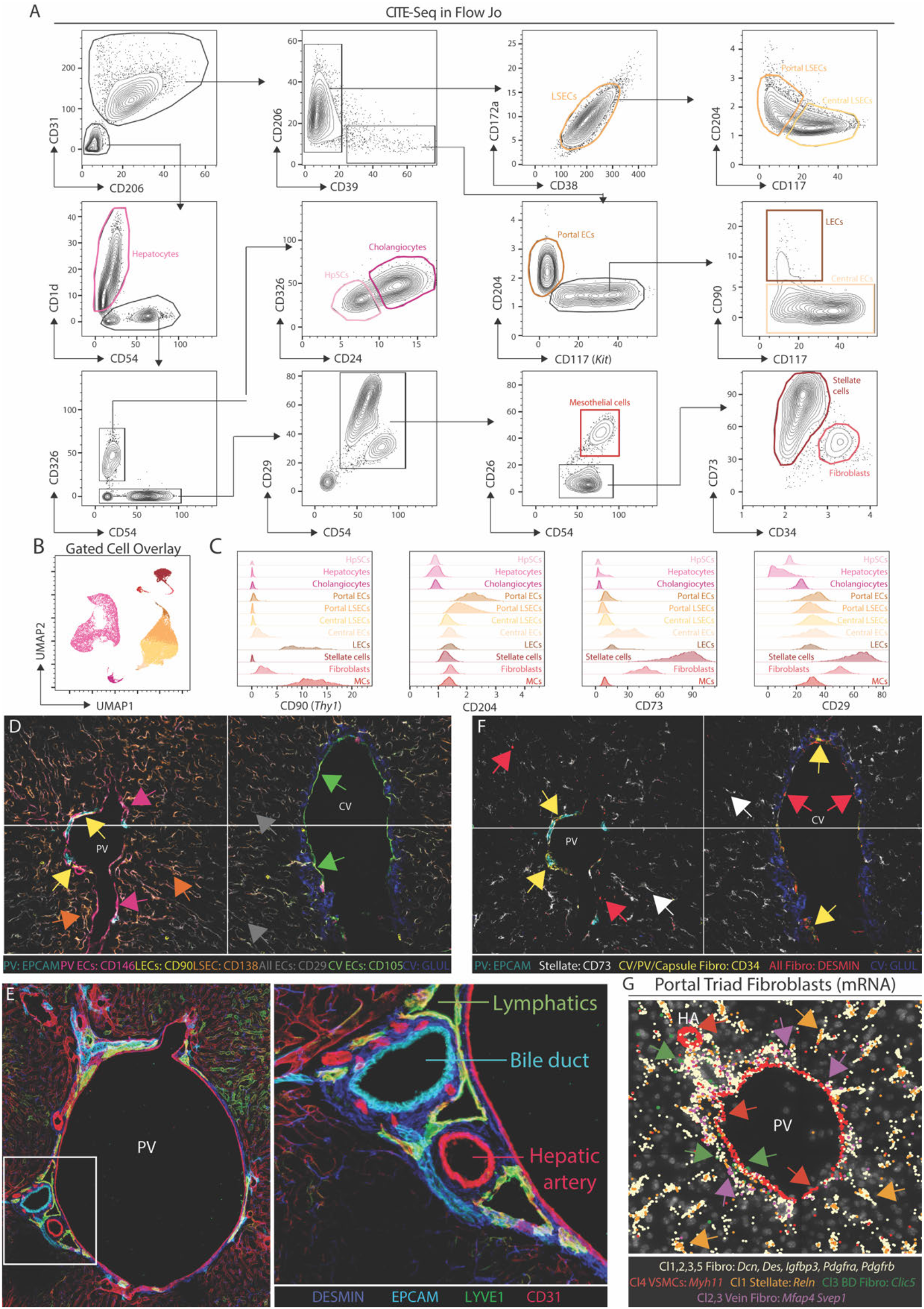
Protein markers of murine CD45^−^ cell subsets. Related to Figure 3. (A) CITE-seq data from the murine CD45^−^ cells in Fig. 3A were exported as an FCS file and an *in-silico* gating strategy identified in FlowJo. (B) Gated cell overlay of populations identified using strategy in A. (C) Expression of CD90, CD204, CD73 and CD29 markers by indicated cell types. (D) Expression of indicated protein markers in 60-plex MICS analysis in endothelial cells. (E) Expression of DESMIN, EPCAM, LYVE1 and CD31 at a portal triad (left) with inset (right). (F) Expression of indicated protein markers in 60-plex MICS analysis in stromal cells. (G) Molecular Cartography of indicated genes and cell types at portal vein. PV; portal vein, CV; central vein. Arrows indicate specific cell types, where color corresponds to cell type/markers. All images are representative of 2-6 mice.

**Fig. S5:**
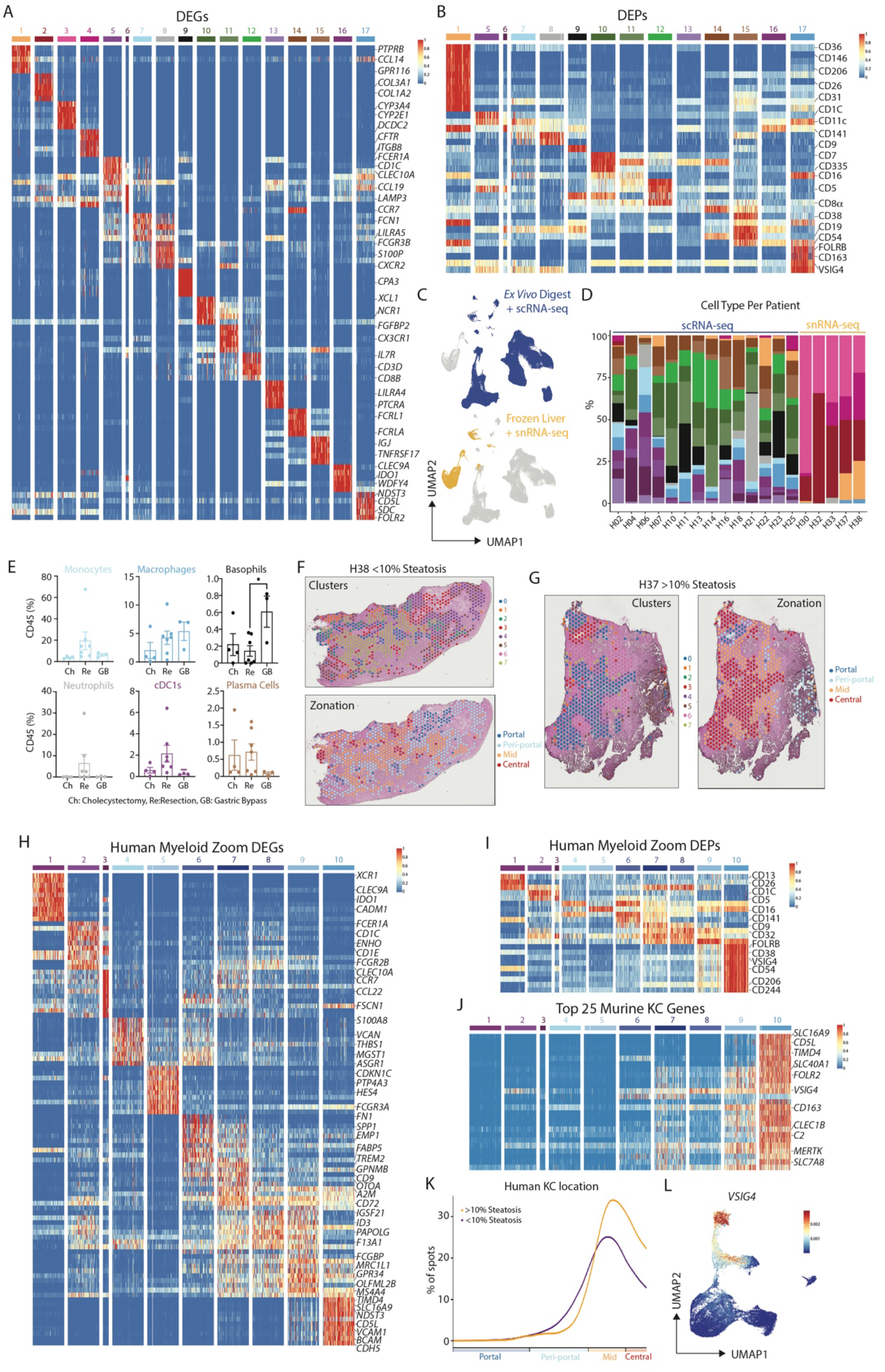
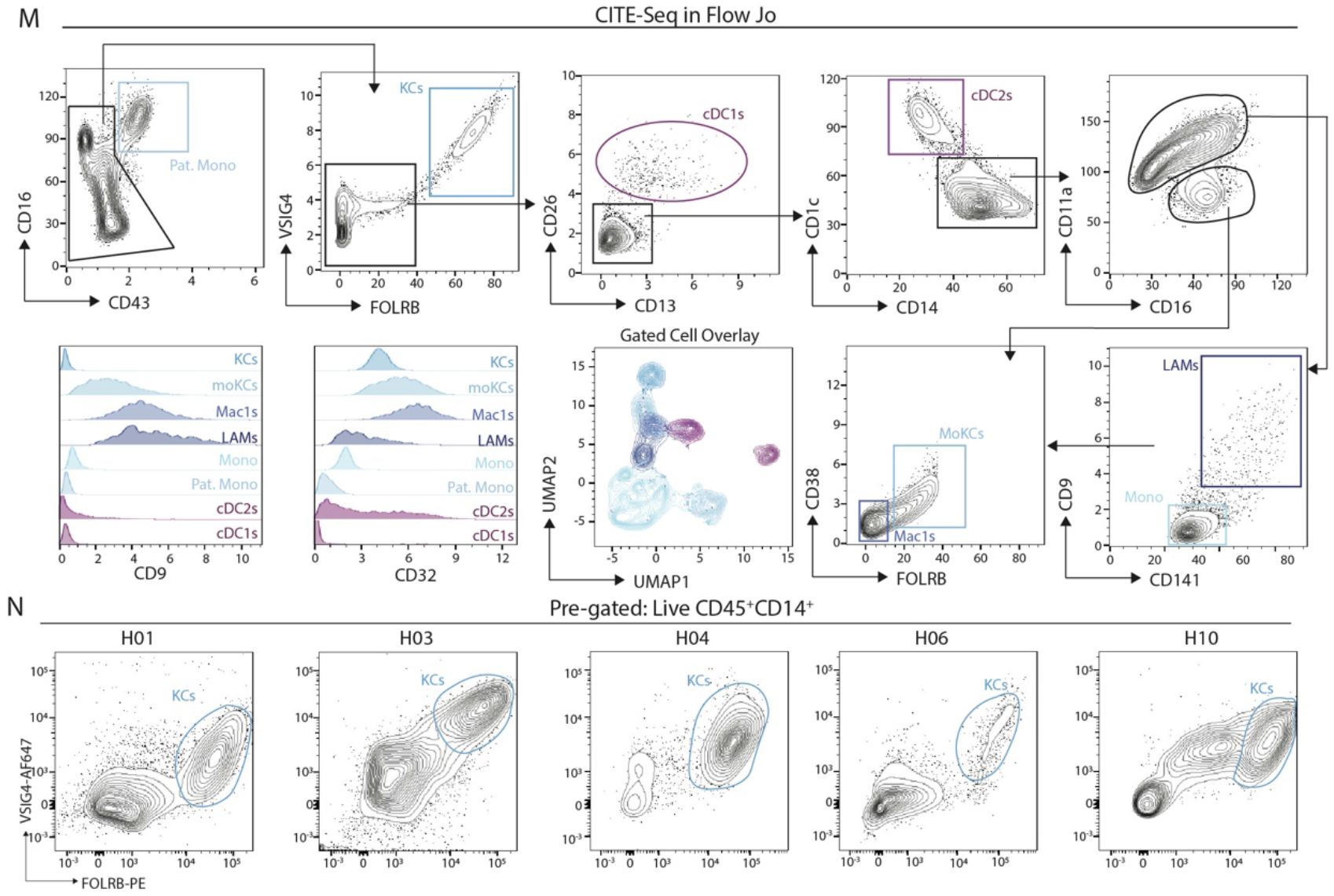
Combination of CITE-seq, scRNA-seq, snRNA-seq and spatial analyses enables generation of a human liver atlas. Related to Figure 4. (A,B) Top DEGs (A) and DEPs (B) for the cell types identified in Fig. 4B. (C) Distinct profiles of cells or nuclei within the UMAP depending on isolation protocol used; 152535 cells from *ex vivo* digestions and 15063 nuclei. (D) Proportion of each cell type per patient profiled. (E) Proportion of indicated cell types as a % of total CD45^+^ cells calculated from *ex vivo* digested samples per surgery type. Ch; Cholecystectomy, Re; Resection, GB; Gastric bypass. *p<0.05 One Way ANOVA with Bonferroni post-test. (F,G) Mapping of Visium UMAP clusters and zonation pattern onto tissue sections from patient H35 (F) and H37 (G). (H,I) Top DEGs (H) and DEPs (I) for cell types identified in Fig. 4H. (J) Top 25 Murine KC genes as expressed by the human myeloid cell clusters. (K) Mapping of KC signature onto Visium trajectory for healthy (purple) and steatotic (orange) livers. (L) Expression of *VSIG4* mRNA within human myeloid cells. (M) *In-silico* gating strategy to isolate distinct myeloid cell populations identified from CITE-seq data. (N) Expression of VSIG4 and FOLR2 by live CD45^+^ cells also expressing CD14 in indicated human liver biopsies by flow cytometry. Data are representative of 21 biopsy samples analyzed.

**Fig. S6:**
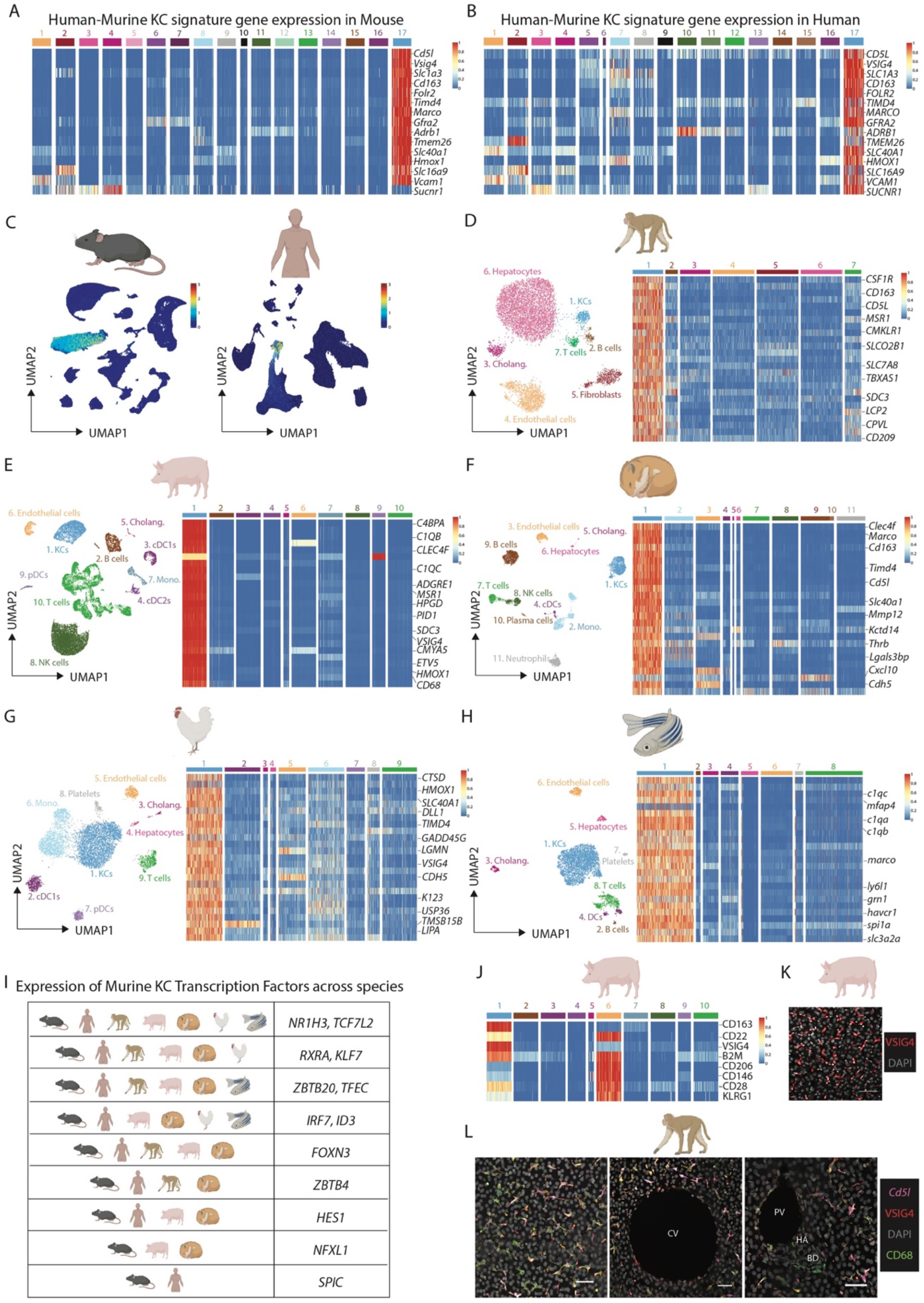
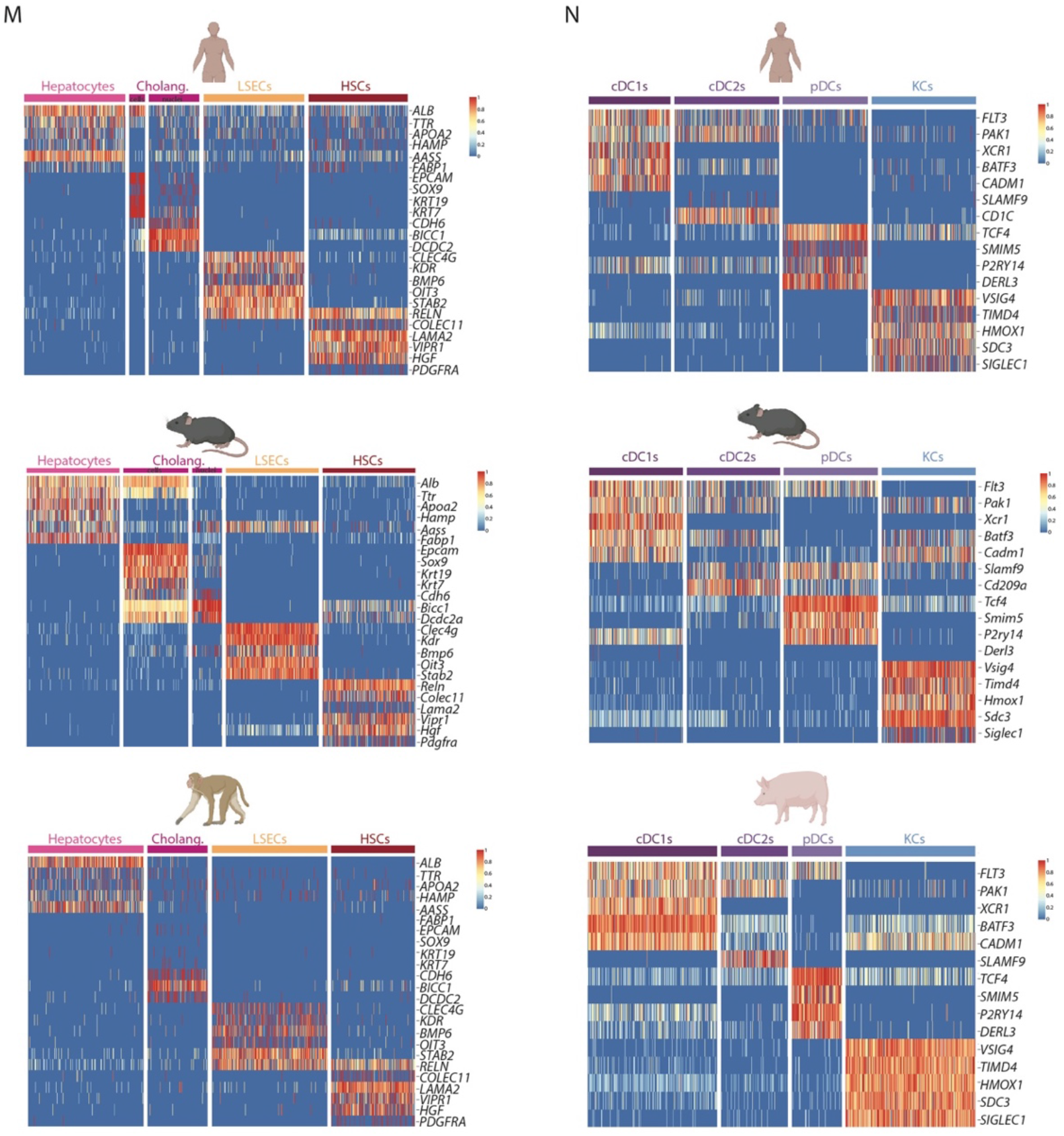
Conserved and unique features of KCs across species. Related to Figure 4. (A,B) Expression of Human-Murine KC signature genes across cell types in mouse (A) and human (B). (C) Unbiased identification of KCs in mouse and human using the human-murine KC signature and the signature finder algorithm (*18*). (D-H) Annotated UMAPs from indicated species and expression of top KC-specific genes compared with other cells per species. (I) Expression of previously identified core murine transcription factors (*8*) by KCs across species. (J) Top DEPs (identified with cross reactive human antibodies) in the pig CITE-seq data. (K) Expression of VSIG4 in the porcine liver by confocal microscopy. (L) Expression of VSIG4, CD68 (protein) and *CD5L* (mRNA) in macaque liver. PV; portal vein, HA; hepatic artery, BD; bile duct. All images are representative of 2 livers. (M,N) Conserved expression of indicated genes across CD45- (M) and CD45+ (N) cell types and species.

**Fig. S7:**
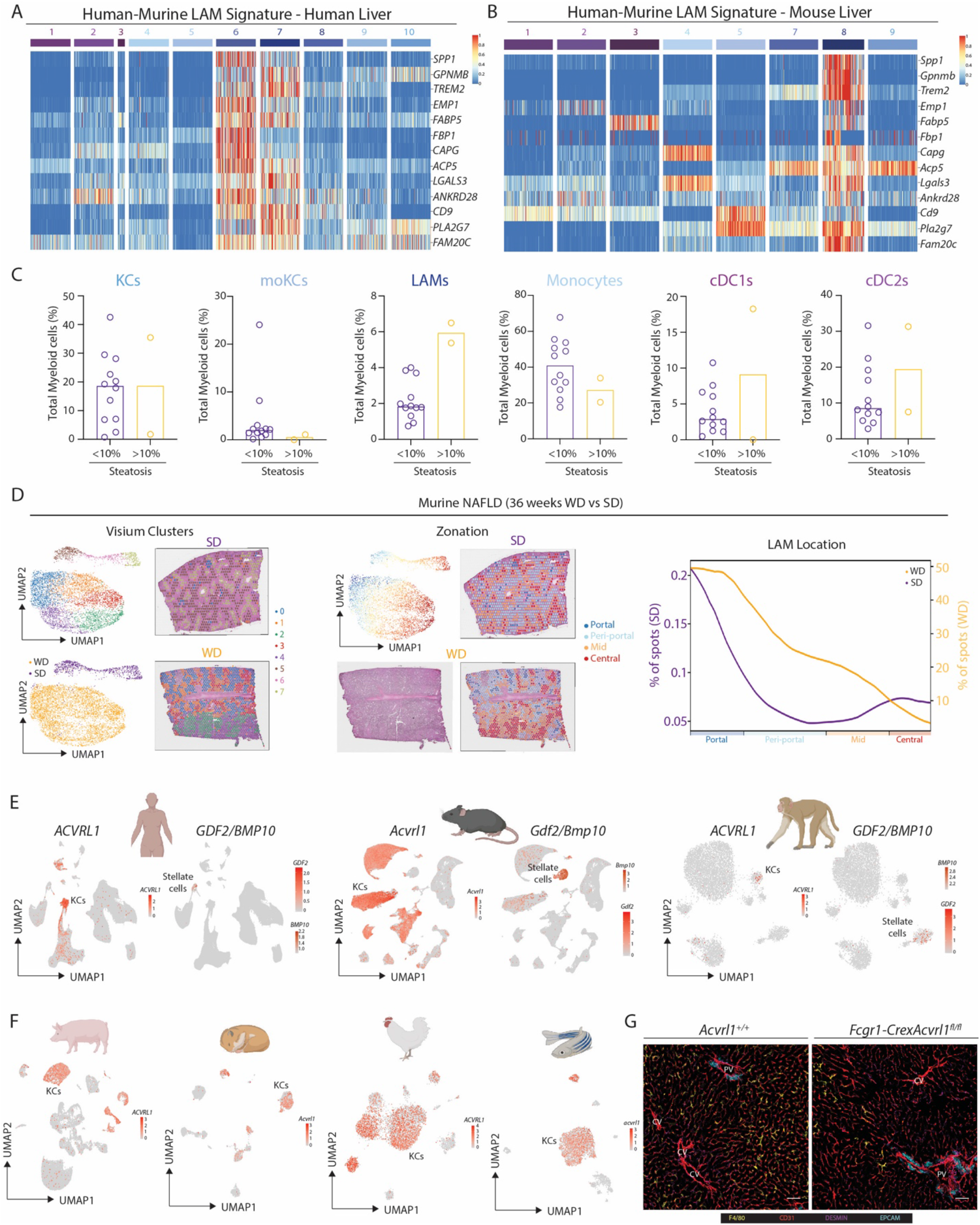
Evolutionarily-conserved signals regulate LAMs and KCs. Related to Figures 5 & 6. (A,B) Expression of conserved Human-Murine bile-duct LAM signature in human (A) and mouse (B) hepatic myeloid cells. (C) Proportion of indicated myeloid cell populations as a % of total myeloid cells in human liver biopsies profiled by scRNA-seq when divided based on presence of steatosis. (D) Mice were fed a Western diet (WD) or Standard diet (SD) for 36 weeks to induce NAFLD and Visium analysis was performed. Analysis is pooled from 1 liver slice from the SD condition and 3 liver slices from the WD condition. Left; cluster and sample annotations, middle; zonation in UMAP and on representative tissue slice and right; location of LAM signature (combination of bile-duct LAM signature from healthy mouse and NAFLD LAM signature from Remmerie et al., 2020) in SD and WD samples along zonation trajectory. (E) Expression of ALK1 (*ACVRL*1), BMP9 (*GDF2*) and *BMP10* in human, mouse and macaque livers where both KCs and stellate cells were profiled. (F) Expression of ALK1 (*ACVRL1, Acvrl1, acvrl1*) in indicated species profiled by scRNA-seq only and thus lacking Stellate cells. (G) Expression of CD31 (ECs), DESMIN (Fibroblasts), F4/80 (Macrophages) and EPCAM (Cholangiocytes) by confocal microscopy in *Fcgr1*-*Cre*x*Acvrl1*^fl/fl^ mice and *Acvrl1*^+/+^ controls. PV; portal vein, CV; central vein. Images are representative of 2 mice per group.

### Supplementary Tables

**Table S1.**

Differentially Expressed Genes (DEG) per cell type in Total Mouse Liver Atlas.

**Table S2.**

Differentially Expressed Proteins (DEP) per cell type in Total Mouse Liver Atlas.

**Table S3.**

Differentially Expressed Genes (DEG) per cell type in Mouse Myeloid cell Atlas.

**Table S4.**

Differentially Expressed Proteins (DEP) per cell type in Mouse Myeloid cell Atlas.

**Table S5.**

Differentially Expressed Genes (DEG) per cell type in Mouse CD45-cell Atlas.

**Table S6.**

Differentially Expressed Proteins (DEP) per cell type in Mouse CD45-cell Atlas.

**Table S7.**

Differentially Expressed Genes (DEG) per cell type in Mouse Stromal cell Atlas.

**Table S8.**

Clinical data from patient liver biopsies used in the study.

**Table S9.**

Differentially Expressed Genes (DEG) per cell type in Total Human Liver Atlas.

**Table S10.**

Differentially Expressed Proteins (DEP) per cell type in Total Human Liver Atlas.

**Table S11.**

Differentially Expressed Genes (DEG) per cell type in Human Myeloid cell Atlas.

**Table S12.**

Differentially Expressed Proteins (DEP) per cell type in Human Myeloid cell Atlas.

**Table S13.**

Differentially Expressed Genes (DEG) per cell type in Total Macaque Liver Atlas.

**Table S14.**

Differentially Expressed Genes (DEG) per cell type in Total Pig Liver Atlas

**Table S15.**

Differentially Expressed Proteins (DEP) per cell type in Total Pig Liver Atlas.

**Table S16.**

Differentially Expressed Genes (DEG) per cell type in Total Hamster Liver Atlas.

**Table S17.**

Differentially Expressed Genes (DEG) per cell type in Total Chicken Liver Atlas.

**Table S18.**

Differentially Expressed Genes (DEG) per cell type in Total Zebrafish Liver Atlas.

**Table S19.**

Differentially Expressed Genes (DEG) per cell type in Human CD45-cell Atlas.

