## Supplementary material for "Spatial proteogenomics reveals distinct and evolutionarily-conserved hepatic macrophage niches": Table S8

| Sample Number Flow | Biopsy used for | Gender | Age | Weight | Height | Reason for hospitalization | Procedure | Concomitant medication | Concomitant disorders/operations | Alcohol intake | Steatosis | Fibrosis |
|---|---|---|---|---|---|---|---|---|---|---|---|---|
| **H01** | Flow Cytometry | male | unknown | unknown | unknown | unknown | unknown | unknown | unknown | unknown | unknown | unknown |
| **H02** | CITE-Seq Flow Cytometry | male | unknown | unknown | unknown | Colorectal cancer with liver metastases | Liver resection | unknown | unknown | unknown | unknown | unknown |
| **H03** | Flow Cytometry | male | unknown | unknown | unknown | Liver resection | Liver resection | unknown | unknown | unknown | unknown | unknown |
| **H04** | scRNA-seq<br>Flow Cytometry | female | 53y | 73kg | 161cm | Symptomatic cholecystolithiasis | Laparoscopic cholecystectomy | Zestril (Lisinopril) 10 mg daily<br>Omeprazol 20 mg daily<br>Lutenyl (Nomegestrol) 5 mg daily<br>Simvastatine 20 mg daily<br>Somatuline (Lanreotide) 90 mg once every 4 weeks<br>D-Cure (Colecalciferol) once every 4 weeks | Pituitary macroadenoma --> acromegaly --> endoscopic hypophysectomy<br>Oophorectomy due to cyst<br>Light form of obstructive sleep apnea<br>Hemorrhoids --> rubber band ligation<br>Diabetes mellitus type 2 --> metformine (not in medication overview..)<br>Pulmonary arterial hypertension<br>Diaphragmatic hernia<br>Stage 2 chronic kidney disease<br>Idiopathic polyneuropathy<br>Cystic hygroma --> surgical removal | 1-2 standard units daily | Mild (10-15%); centrilobular; macrosteatosis | No |
| **H05** | Flow Cytometry | male | 56y | 79kg | 176cm | Colorectal cancer with synchronous multiple liver metastases | Parenchyma-sparing open liver resection | In advance: 4 cycles of FOLFOX (Leucovorin/ Fluorouracil/Oxaliplatin) + panitumumab<br>Cortisone ointment due to chemotherapy-induced rash | Low back pain<br>Cervical spondylosis | Unknown | Mild (10-15%) | No |
| **H06** | CITE-seq Flow Cytometry | male | 72y | 87kg | 174cm | Relapse of colorectal liver metastases (segment VI/VII) | Open liver resection | In advance: 8 cycles of CAPOX (Capecitabine/Oxaliplatin) | COPD (GOLD II)<br>Arrhythmia<br>Prostate cancer (Gleason 6)<br>Colorectal cancer with liver metastases --> laparoscopic sigmoid colectomy + microwave ablation of liver lesions<br>Squamous cell carcinoma --> chemoradiotherapy with Cisplatin/Etoposide | Unknown | No | No |
| **H07** | scRNA-Seq Flow Cytometry | female | 46y | 69kg | 160cm | Hepatocellular adenoma (segment III) | Liver resection with adhesiolysis | Zaldiar (Tramadol/Paracetamol) 37.5/325 mg daily<br>Livial (Tibolon) 2.5 mg daily<br>Citalopram 20 mg daily<br>Nortrilen (Nortriptyline) 25 mg daily | Endometriosis --> hysterectomy<br>Oophorectomy of both ovaries<br>Appendectomy<br>Bilateral hydrosalpinx --> salpingectomy<br>Laparoscopic caecectomy<br>Colpopromontoriopexy<br>Total colectomy with ileorectal anastomosis<br>Hypovitaminosis D --> startup of supplements (not in medication overview..) | Unknown | No | No |
| **H08** | Flow Cytometry | male | 73y | 80kg | 180cm | Symptomatic cholelithiasis | Laparoscopic cholecystectomy | Loortan (Losartan) 50 mg daily<br>Relvar (Fluticason/Vilanterol) 92/22 µg daily<br>Ventolin (Salbutamol) | Asthma<br>Pulmonary arterial hypertension<br>Granulomatous rosacea<br>Surgical removal of 6 tubular adenomas with low-grade dysplasia | Social drinker | No | No |
| **H09** | Flow Cytometry | male | 37y | 90kg | 188cm | Colorectal cancer with solitary liver metastasis (segment V) | Laparoscopic resection of | 4 cycles of FOLFIRINOX (Leucovorin/ Fluorouracil/Irinotecan/Oxaliplatin) bevacizumab | | Unknown | No | No |
| **H10** | scRNA-seq<br>Flow Cytometry<br>Confocal Microscopy | male | 55y | 116kg | 173cm | Morbid obesity | Laparoscopic gastric bypass | Selozok (Metoprolol) 100 mg daily<br>Zestril (Lisinopril) 20 mg daily<br>Asaflow (Acetylsalicylic acid) 80 mg daily<br>Zocor (Simvastatine) 40 mg daily<br>CetiSandoz (Cetirizine) 10 mg daily | Dermatographic urticaria<br>Tonsillectomy<br>Acute myocardial infarction complicated by major arrhythmias --> percutaneous coronary intervention<br>Pulmonary arterial hypertension<br>Hyperlipidemia<br>Intersphincteric abscess | 4 standard units daily | No | No |
| **H11** | CITE-Seq Flow Cytometry | female | 53y | 114kg | 167cm | Morbid obesity | Laparoscopic gastric bypass | L-Thyroxine 175 µg on odd days<br>L-Thyroxine 150 µg on even days<br>Omeprazol 40 mg twice daily<br>Simvastatine 20 mg daily<br>Rivotril (Clonazepam) 2 mg if necessary<br>Mirtazipine 15 mg daily<br>Tramium (Tramadol) 100 mg daily<br>Temesta (Lorazepam) 2,5 mg if necessary<br>Daflon (Diosmine/Flavonoids) 450/50 mg if necessary | Laparoscopic sacropexy<br>Hashimoto's thyroiditis<br>Spinal stenosis --> laminectomy<br>Carpal tunnel release<br>Hepatic steatosis<br>Insomnia<br>Reflux<br>Tremor | 1 standard unit monthly | Mild (20%) | Clear pericellular fibrosis |
| **H12** | Flow Cytometry | male | 48y | 110kg | 181cm | Symptomatic cholecystolithiasis | Laparoscopic cholecystectomy | Paroxetine 20 mg daily<br>Pantomed (Pantoprazol) 40 mg twice daily<br>Spasmomen (Otilonium bromide) 40 mg three times a day | Renal colic<br>Gastro-esophageal reflux disease (Grade A)<br>Several abdominal and dorsal lipomas --> resection<br>Laparoscopic appendectomy with peritonitis | 3 standard units weekly | 5% steatosis | Mild pericellular fibrosis |
| **H13** | CITE-seq Flow Cytometry | male | 73y | 107kg | 172cm | Symptomatic cholecystolithiasis | Laparoscopic cholecystectomy | Asaflow (Acetylsalicylic acid) 80 mg daily<br>Bisoprolol 2.5 mg daily<br>D-cure (Colecalciferol) once weekly<br>Efexor (Venlafaxine) 75 mg daily<br>Glucophage (Metformin) 500 mg twice daily<br>Humalog (Insulin lispro) 10 IU three times daily<br>Lantus (Insulin glargine) 40 IU once daily<br>L-Thyroxine 125 µg daily<br>Lisinopril 5 mg daily<br>Simvastatine 40 mg daily<br>Paracetamol 1 g if necessary | Diabetes mellitus type 2 with micro- and macrovascular complications<br>Bursitis left elbow --> bursectomy<br>Cataract --> surgery<br>Autonomous struma --> total thyroidectomy<br>Syncope with hypoglycemia and hypotension --> hospitalization<br>Chronic kidney disease<br>Prostatitis --> hospitalization<br>Spinal stenosis --> surgery | Unknown | 5% steatosis | Clear pericellular fibrosis |
| **H14** | scRNA-seq<br>Confocal Microscopy<br>Flow Cytometry | female | 75y | 63kg | 160cm | Symptomatic cholecystolithiasis | Laparoscopic cholecystectomy | Cholecalciferol 10 µg daily<br>Domperidon 10 mg if necessary | Carpal tunnel release<br>Disturbed glycemia<br>Hypercholesterolemia<br>Cataract --> surgery | Unknown | >5% steatosis | Very mild pericellular fibrosis |
| **H16** | CITE-seq<br>Confocal Microscopy<br>Flow Cytometry | female | 66y | 56kg | 156cm | Colorectal cancer with liver metastases (segment II/VIII) | Liver resection with adhesiolysis | In advance: 4 cycles of FOLFOX (Leucovorin/ Fluorouracil/Oxaliplatin) | Wertheim-Meigs operation (in 1998) | Unknown | <5% steatosis | No |
| **H17** | Flow Cytometry | female | 33y | 64kg | 178cm | Symptomatic cholecystolithiasis | Laparoscopic cholecystectomy | Paracetamol 1 g if necessary<br>Bellina (Ethinylestradiol/Chlormadinone acetate) 0.03/2 mg daily | Endometriosis | Unknown | No | Mild pericellular fibrosis |
| **H18** | CITE-Seq Flow Cytometry | female | 28y | 52kg | 157cm | Hepatocellular adenoma (segment V/VI) | Laparoscopic liver resection + cholecystectomy | Movicol (Macrogol + electrolytes) if necessary<br>Paracetamol 1 g if necessary<br>Tradonal (Tramadol) 50 mg if necessary | | Unknown | No | Mild pericellular fibrosis |
| **H19** | Flow Cytometry | female | 25y | 67kg | 160cm | Colorectal cancer with synchronous diffuse liver metastases | Liver resection + microwave ablations | Clexane (Enoxaparin sodium) 40 mg daily<br>Lindynette 30 (Ethinylestradiol/Gestodene) 0.03/0.075 mg daily<br>Magnesium daily<br>Minocycline 100 mg daily<br>Mometasone nasal spray 50 µg twice daily<br>Pantomed (Pantoprazol) 40 mg daily<br>Paracetamol 1 g if necessary<br>Tradonal (Tramadol) 50 mg if necessary<br>In advance: 8 cycles of FOLFOX (Leucovorin/ Fluorouracil/Oxaliplatin) + panitumumab | Acne<br>Asthma<br>Hay fever | Unknown | No | No |
| **H20** | Flow Cytometry | female | 51y | 80kg | 173cm | Colorectal cancer with metachronous liver metastases | Liver resection | In the past: adjuvant FOLFOX (Leucovorin/ Fluorouracil/Oxaliplatin)<br>In advance: 8 cycles of FOLFIRINOX (Leucovorin/ Fluorouracil/Irinotecan/Oxaliplatin) + bevacizumab | | Unknown | 5-10% steatosis | Pericellular fibrosis |
| | | | | | | | | Amlodipine 5 mg daily<br>Betahistine 16 mg three times daily<br>Mirabegron 25 mg daily | Diabetes mellitus type 2 with peripheral polyneuropathy<br>Metabolic syndrome<br>Total hysterectomy | | | |

| | | | | | | | | | | | | |
|---|---|---|---|---|---|---|---|---|---|---|---|---|
| **H21** | scRNA-seq<br>Flow Cytometry | female | 77y | 68kg | 156cm | Hepatocellular carcinoma (segment II/VII) | Liver resection | Acetylsalicylic acid 100 mg daily<br>Clexane (Enoxaparin sodium) 40 mg daily<br>Duloxetine 60 mg daily<br>Deanxit (10 mg melitracen + 0.5 mg flupentixol) once daily<br>Metformin 500 mg daily<br>Lormetazepam 1 mg daily<br>Oxycodon 20 mg daily<br>Pantoprazol 40 mg daily<br>Paracetamol 1 g four times daily<br>Prolopa (100 mg levodopa + 25 mg benserazide) three times daily<br>Magnesium 3g daily<br>Simvastatine 40 mg daily | Cholecystectomy<br>Diverticulosis<br>Hemorrhoids<br>Gastric sleeve<br>Parkinson<br>Cystadenocarcinoma --> bilateral overiectomy and appendectomy + chemotherapy (Endoxan, Platinol) | Unknown | <5% steatosis | Cirrhosis |
| **H22** | scRNA-seq<br>Flow Cytometry | female | 55y | 66kg | 169cm | Liver injury (segment VI) | Laparoscopic liver resection | Unknown | Migraine<br>Meningitis | Unknown | No | No |
| **H23** | scRNA-seq<br>Flow Cytometry | female | 75y | 142kg | 168cm | Morbid obesity | Roux en-Y gastric bypass | Unknown | hypothyroidism<br>appendectomy | Unknown | No | 5-10% fibrosis |
| **H24** | Flow Cytometry | male | 58y | 74kg | 177cm | Metastasis of colorectal adenocarcinoma (segment VIII) | Liver resection | Atorvastatine (Lipitor) 40 mg daily<br>Minocycline 100 mg daily<br>Paracetamol 1 g four times daily if necessary<br>Tradonal (Tramadol) 50 mg four times daily if necessary | Asymptomatic chronic calcifying pancreatitis<br>Nefrolithiasis<br>Lumbar hernia | Sporadic | 5-10% steatosis | No |
| **H25** | scRNA-seq<br>Confocal +RNAScope<br>Flow Cytometry | female | 49y | 99kg | 176cm | Symptomatic cholecystolithiasis | Laparoscopic cholecystectomy | Seretide (50 µg salmeterol + 250 µg fluticason) one dose daily<br>Ventolin (Salbutamol)<br>Nasonex (mometason) 2 times twice daily<br>Flixonase (fluticason)<br>Microgynon (30 µg ethinylestradiol + 150 µg levonorgestrel) once daily | Asthma<br>Nasal polyposis grade 3<br>Obesity | No | 5-10% steatosis | Minimal pericellular fibrosis |
| **H26** | Confocal + RNAScope | male | 57y | 120kg | 191cm | Colorectal cancer with liver metastases | Liver resection | In advance: 4 cycles of CAPOX (Capecitabine/Oxaliplatin)<br>Metoprolol<br>Enalapril<br>Allopurinol | Arrhythmia<br>Sinusitis<br>Arterial hypertension<br>Gout | Unknown | 20% steatosis | Mild pericellular fibrosis |
| **H29** | Confocal + RNAScope | female | 67y | 106kg | 164cm | Morbid obesity | Gastric bypass | Nobiten (nebivolol) 2.5 mg twice daily<br>Entresto (49 mg sacubitril + 51 mg valsartan)<br>Procoralan (ivabradine) 5 mg<br>Pantomed (pantoprazol) 40 mg once daily<br>Cymbalta (duloxetine) 60 mg once daily<br>Zolpidem 10 mg once daily<br>Calcium<br>Mirtazipine 15 mg daily<br>Spironolactone 25 mg daily<br>Montelukast once daily<br>Relvar (Fluticason/Vilanterol) 184/22 µg daily<br>Incruse (umeclidinium) once daily<br>Duovent (ipratropium + fenoterol) if necessary<br>Azithromycine 250 mg three times daily<br>Mometasone nasal spray (50 µg) 2 doses once daily | Osteopenia<br>Hemorrhoids<br>Hysterectomy/ovariectomy<br>Cholecystectomy<br>Gastritis<br>Cardiomyopathy<br>Asthma with persistent bronchial obstruction<br>Depression | Unknown | 10% steatosis | Minimal pericellular fibrosis |
| **H30** | snRNA-seq | female | 64y | 96kg | 159cm | Obesity class 2 | Gastrc Bypass | Asaflow (Acetylsalicylic acid) 80 mg daily | Appendectomy<br>Cholelithiasis --> cholecystectomy<br>Varicectomy | No | 15% | No |
| **H32** | snRNA-seq | male | 33y | 123kg | 174cm | Obesity class 3 | Gastrc Bypass | / | / | Unknown | 50% | No |
| **H33** | snRNA-seq | male | 67y | 82.7kg | Unknown | Liver abscess | Resection | Lisinopril 5 mg daily<br>Asaflow (Acetylsalicylic acid) 80 mg daily<br>Paracetamol 1 g if necessary<br>Tradonal (Tramadol) 50 mg if necessary<br>Tresiba (Insulin degludec) 16 IU once daily<br>Novorapid (Insulin aspart) 10 IU three times daily | Cholecystectomy<br>Diabetes mellitus type 1 | Unknown | No | Mild pericellular fibrosis |
| **H35** | Visium | female | 59y | 102kg | 159cm | Obesity class 3 | Gastric Bypass | Co-Bisoprolol (bisoprolol/hydrochlorothiazide) 5/12.5 mg daily<br>Pantoprazol 40 mg daily | Tonsillectomy<br>Adenotomy<br>Appendectomy<br>Nefrolithiasis<br>Menopause<br>Arterial hypertension | Unknown | 35-40% | No |
| **H36** | Visium | male | 59y | 55kg | 175cm | Colorectal cancer with liver metastases | Resection | Epipen (epinefrine) 0.3 mg if necessary | Food allergy | 6 standard units daily | No | mild pericellular & periportal |
| **H37** | snRNA-seq<br>Visium | male | 58y | 118kg | 176cm | Obesity class 2 | Gastric Bypass | Asaflow (Acetylsalicylic acid) 80 mg daily<br>Olmetec (olmesartan/medoxomil) 40/25 mg daily<br>Bisoprolol 10 mg daily<br>Metformin 850 mg three times daily<br>Pantoprazol 40 mg daily<br>Atorvastatine 40 mg daily<br>Nortrilen (Nortriptyline) 25 mg three times daily<br>Baclofen 5 mg three times daily<br>Paracetamol 1 g three times daily<br>Pregabaline 300 mg twice daily<br>Sulpiride 50 mg twice daily | Diabetus mellitus type 2<br>Obstructive sleep apnea<br>Depression<br>Coronary artery disease<br>Arterial hypertension<br>Hypercholesterolemia | Unknown | 70% | Clear pericellular fibrosis |
| **H38** | snRNA-seq<br>Visium | female | 68y | 95kg | 174cm | Symptomatic cholecystolithiasis | Cholecystectomy | / | Obesity | Unknown | 10% | Mild pericellular fibrosis |
